# Interferon-induced lysosomal membrane permeabilization and death cause cDC1-deserts in tumors

**DOI:** 10.1101/2022.03.14.484263

**Authors:** E. Aerakis, A. Chatzigeorgiou, M. Alvanou, M. Matthaiakaki-Panagiotaki, I. Angelidis, D. Koumadorakis, A. Galaras, P. Hatzis, D. Kerdidani, M. Makridakis, A. Vlachou, B. Malissen, S. Henri, M. Merad, M. Tsoumakidou

## Abstract

T cell immunity requires antigen capture by conventional dendritic cells (cDCs), digestion and transfer to draining lymph nodes for presentation to antigen-inexperienced T cells. cDCs type I excel as cancer-antigen presenting cells, due to their ability to phagocytose, slowly digest apoptotic cancer cells and translocate cancer antigens to the cytosol for loading to MHCI and cross-presentation to CD8 T cells ^1–3^. In tumor tissues cDCs1 become particularly scarce and this restricts anti-tumour immunity, immunotherapy responses and patient survival ^4–8^. Tumor cDC1 paucity is not fully understood and no specific treatment currently exists. Here, we find that type I interferons (IFN) induce lysosomal stress, lysosomal membrane permeabilization (LMP) and lysosomal-dependent cell death (LDCD) in cDCs1. Two parallel pathways downstream of IFNAR1 converged to induce cDC1 LDCD. Up-regulation of expression of lysosomal genes enhanced the proteolytic activity of lysosomes, while IFN-inducible guanylate binding protein-2 (GBP-2) accumulated in the membrane of the stressed lysosomes, leading to LMP, proteolytic enzyme release and death. Protease inhibition or GBP-2 repression rescued cDCs1 from LDCD and boosted their anti-tumor efficacy. GBPs are amongst the most abundant IFN-induced genes and known to form toxic pores in pathogen-containing vacuoles and pathogen membranes ^9^. GBP-2-driven LMP is likely due to the ability of GBP-2 to form pores on the lysosomes of cDC1s. This might have evolved as a physiological mechanism of antigen translocation to the cytosol for cross-presentation ^10^. We anticipate our findings to be a starting point for more rational cDC1-directed immunotherapies. For instance, protease inhibition, GBP-2 downregulation or induced expression of LMP repair machinery may boost cDC1 efficacy in adoptive cell therapies or their use as live vaccines^11–13^.

## Introduction

“All things excellent are as difficult as they are rare”, Baruch Spinoza.

Immune cell co-evolution led to the emergence of dendritic cell (DC) subsets with contextual specifications in cancer immunity ^14–16^. Conventional DCs type 1 (cDCs1) surpass other DCs in their ability to prime antigen-inexperienced cancer specific CD8 T cells and are regarded as the most potent anti-tumour APCs ^17^. Prolonged survival is observed in cancer patients with higher cDC1 infiltration ^4,7,18,19^. cDC1 ablation or editing in animal models compromises immunity and accelerates cancer progression ^20–22^. Cancer immunotherapy responses depend on cDC1 mobilization ^5,18,23–25^. The non-redundant anti-tumour functions of cDCs1 are particularly impressive considering that they are rare in tissues and become extremely scarce in solid tumours ^6,8,26^.

The enhanced cross-priming capacity of cDCs1 is attributed to their superiority in phagocytosing cancer cell debris via DNRG1 (also termed CLEC9A) and in slowly degrading antigens for long-term antigen storage and cross-presentation ^1–4,20,27,28^. cDCs1 largely depend on the cytosolic pathway of cross-presentation, during which antigens are released from the phagosome to the cytosol, degraded by the proteasome and loaded on MHCI ^29^. DNRG1 activates phagosomal rupture and cancer antigen release to the cytosol before fusion with the lysosome ^1^. cDCs1, but not other DC subsets, can also release antigens to the cytosol after phagosome-lysosome fusion ^30^. This process should be very tightly regulated to prevent uncontrolled release of toxic proteases that can cause irreparable cell damage and trigger a type of death that is known as lysosome-dependent cell death (LDCD) ^13,31^. Controlled antigen translocation in the late cytosolic pathway might involve the Endoplasmic Reticulum-Associated Protein Degradation (ERAD) member Sec61 or a yet unknown transmembrane pore complex ^10,32^.

cDC1 paucity in tumor is not fully understood. Tumor-produced granulocyte-stimulating factor downregulates bone marrow cDC progenitors, but this should not impact specifically cDCs1 ^33^. Decrease in cDC1 chemokines impedes tumor cDC1 infiltration, but recruitment failure cannot explain extinction ^25,34,35^. Adding another level of complexity, tumor-draining LNs themselves become deserted from cDCs1 as cancer progresses and it has been postulated that this is due to impaired LN migration ^36^. Here, we find that type I interferons (IFN) induce lysosomal stress, lysosomal membrane permeabilization (LMP) and LDCD in cDC1s. Two parallel pathways downstream of IFNAR1 converged to induce LDCD. Up-regulation of lysosomal gene expression enhanced lysosome proteolytic activity, while IFN-inducible guanylate binding protein-2 (GBP-2) accumulated in the membrane of the lysosomes, leading to LMP, proteolytic enzyme release and death. Protease inhibition or GBP-2 repression rescued cDC1s from LDCD and boosted their anti-tumor efficacy. Current treatment strategies to increase numbers of cDCs1 in cancer patients are based on exogenous replenishment, mobilization of bone-marrow progenitors or intratumoral injection of cDC1-attracting chemokines ^11^. We anticipate our findings to be a starting point for more rational cDC1-directed immunotherapies. For instance, protease inhibition, GBP-2 repression or induced expression of LMP repair machinery may boost cDCs1 efficacy in adoptive cell therapies or their use as live vaccines ^11–13^.

## RESULTS

### cDC1 lysosomes enter a lysosomal stress state in tumours

To decode the complexity of tumor-induced cDCs1 we leveraged orthotopic lung cancer models developed by direct injection of lung cancer cells in the left lung lobe of syngeneic mice. These mice develop solitary lung tumors, allowing tumor excision and the study of pure tumor-infiltrating cells ^37,38^. First, we set out to investigate whether anti-tumour immunity was dependent on cDCs1 in our model. We (Malissen and colleagues) recently developed a cDC1 specific XCR1^cre^ line in which a Cre recombinase and a fluorescent reporter are co-expressed under the control of the Xcr1 gene, in a manner that maintains XCR1 expression ^28^. We crossed the Xcr1cre line with a loxP-STOP-loxP DTA line to constitutively ablate cDCs1 and studied two lung adenocarcinoma cell lines of different aggressiveness and immunogenicity. The commercially available LLC cell line gives rise to lung tumors that are poorly immunogenic and grow fast, with mice succumbing at ∼ 2 weeks, while the autochthonous urethane-induced lung cancer cell line CULA gives rise to lung tumors that are highly immunogenic and grow slowly, with mice succumbing at ∼ 4 weeks (**Suppl. Fig. 1**). To quantify cancer antigen specific responses and tumor burden both lines were transduced with fluorescent ovalbumin viral vectors. XCR1^cre^LSL-DTA mice developed larger tumors compared to control mice, an effect that was more prominent with the CULA line (**Suppl. Fig 2**). FACS analysis confirmed the successful and specific ablation of intratumoural cDCs1 and revealed a parallel decrease in intratumoral CD4 and CD8 T cells -again more prominent in the CULA line (**Suppl. Fig 2**). Likewise, staining with tetramers that recognize ovalbumin-specific CD8 T cells, indicated that cDC1 ablation impacts cancer-specific T cytotoxic cells (**Suppl. Fig. 2**). We conclude that our lung cancer models are cDC1-dependent and can thus be safely used to study tumor-driven cDC1 states.

Due to their scarcity in scRNAseq datasets, tumour cDCs1 have not been systematically compared to their healthy tissue counterparts ^19,36,39–41^. Single-cell methods inherently suffer from limitations in the recovery of complete transcriptomes due to the prevalence of transcriptional dropout events. To get a focused high-resolution view of the cDC1 transcriptome we analyzed by bulk RNAseq cDCs1 purified from LLC and CULA lung tumors versus healthy lungs **(Suppl. Fig. 3)**. To avoid contamination by the mReg cDC subset which may express low levels of XCR1, we gated on XCR1^+^CD11b^-^ cDCs. A total of 9.720 genes were identified. Principal Components Analysis (PCA) showed that tumor and healthy cDCs1 separate in two distinct clusters **(Fig. 1a)**. One outlier sample was excluded from further analysis. Surprisingly, differential expression analysis found only 30 genes that were highly upregulated in tumor derived cDCs1 and 10 that were highly downregulated, in comparison with healthy cDC1s. Among top downregulated were genes involved in cell structure and adhesion (*Pdlim1, Ahnak, CD44, Serpinb8*), while among top upregulated was the well-known inhibitory molecule PDL1, interferon-stimulated genes (*Gbps2, Gbp5, Gbp7, Stat1 and Irf1*) and several genes involved in lysosomal processes. Specifically, components of the lysosomal membrane that mediate lysosomal trafficking and fusion with other membrane-bound organelles and the plasma membrane (*CD63, Gga2 and Gpnmb*) and the lysosomal proteolytic enzymes *Ctsd* and *Lgmn* (**Fig. 1a**). LLC tumor cDCs1 showed a more robust tumor associated profile compared to CULA tumor cDCs1. We hypothesize that this is due to the higher aggressiveness of LLC tumors. Likewise, upregulated genes analyzed by DAVID showed significant enrichment in pathway activities related to IFNs and lysosomes in cDC1s from tumors (**Fig. 1a**). To validate our findings, we re-analyzed our publicly available scRNAseq dataset (Merad and colleagues) of Kras^G12D^P53^-/-^ (KP) lung tumours and healthy lungs ^42^. We filtered out from our analysis cDCs2, mRegs and contaminants and focused on the cDC1 cluster. Two large (C1, C2) and two small cDC1 sub-clusters (C3, C4) were identified by uniform manifold projection (UMAP). The C1 sub-cluster was almost exclusively derived from the tumor-bearing lungs, while C2 from healthy lungs. (**Fig. 1b**). Functional enrichment analysis of upregulated genes performed by DAVID again showed significant enrichment in lysosomal processes and type I IFN pathways in C1 (**Fig. 1b, Suppl. Fig. 4**). Accordingly, top enriched GO terms/KEGG pathways were highly concordant between bulk and scRNAseq datasets (**Fig. 1c**). Collectively, two independent datasets across 3 different lung cancer models suggest that cDCs1 become transcriptionally biased towards lysosome active states in tumours.

**Figure 1.**
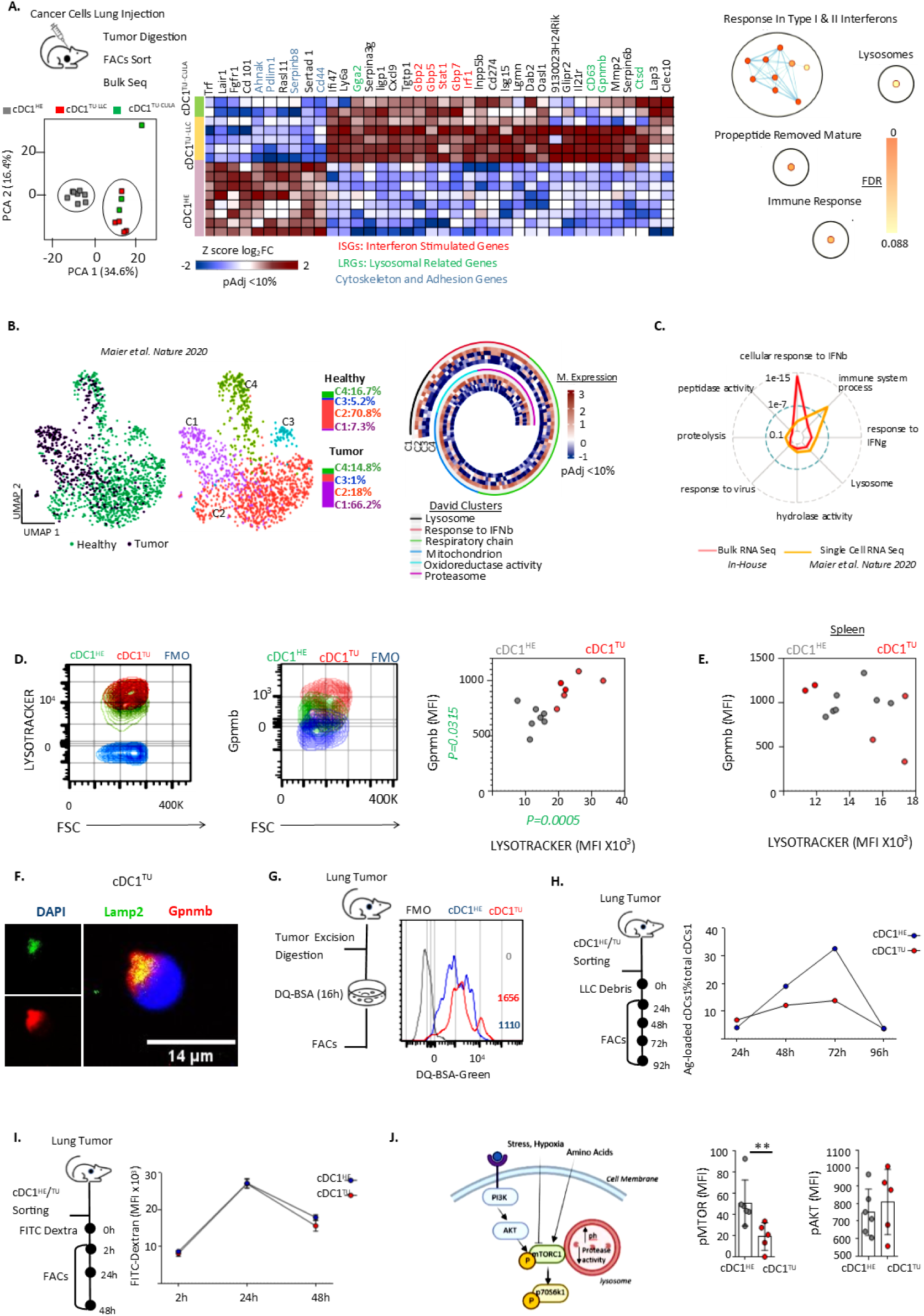
Primary cDCs1 enter a lysosomal stress state in lung tumours. **A**. The LLC or the autochthonous CULA lung cancer cell line were inoculated in the left lung lobe of mice. LLC lung tumors (TU-LLC), CULA lung tumours (TU-CULA) and control healthy lungs (HE) were excised. Lin^-^CD11c^+^MHCII^+^CD11b^-^XCR1^high^ cDCs1 were FACS-sorted and analyzed by bulk RNAseq. Left to right. PCA plot of biological replicates. Heatmap of all differentially expressed genes. DAVID enrichment analysis of overexpressed genes in cDC1^TU-LLC^ and cDC1^TU-CULA^. **B.** Metastatic KP lung tumor and healthy lung cDC1 transcriptomes were extracted from a public murine scRNAseq dataset ^42^. Left to right. UMAP color-coded based on tissue of origin and clusters. Heatmap depicts up-regulated genes in KP lung tumor cDCs1 that were found enriched in pathway enrichment analysis by DAVID. **C.** Common enriched pathways between bulk RNAseq and scRNAseq datasets. **D.** LLC tumours and healthy lungs were excised, stained with an acidotropic probe (lysotracker) and the lysosomal stress marker Gpnmb (intracellular) and analyzed by FACS, gating on cDC1^TU^ and cDC1^HE^. **E.** As in D, but this time spleens were excised and analyzed by FACS. **F.** Co-localization of Gpnmb and lysosomal membrane protein LAMP-2 in cDC1^TU^. **G.** 24h culture of cDC1^TU^ and cDC1^HE^ with fluorogenic substrate for proteases (DQ-BSA) and FACS. **H.** 4-day culture of cDC1^TU^ and cDC1^HE^ with apoptotic fluorescent LLC cells and FACS. **I.** 2-day culture of cDC1^TU^ and cDC1^HE^ with the endocytic probe Dextran and FACS. **J.** Left to Right. Graphical summary of the mTORC1 pathway. Active mTORC1 and AKT quantified by phospho-FACS in cDC1^TU^ and cDC1^HE^. **D-J**. Representative or cumulative data from at least 2 independent experiments with 4-6 biological replicates per group per experiment. Each biological replicate of FACS-sorted cDCs1 was pooled from 3-5 mice. Error bars, mean± sem; **p<0.01 two-tailed unpaired t-test

A key component of lysosomal activities is its pH. Thus, we performed a fluorescent pH reporter assay using a dye (LysoTracker) that stains cellular acidic compartments. We also quantified intracellular Gpnmb (Glycoprotein Non-metastatic Protein B). Gpnmb concentrates on the membrane of dysfunctional lysosomes in lysosomal storage disorders ^43,44^. It is considered a marker of lysosomal stress ^43,44^. Higher Lysotracker fluorescence and intracellular Gpnmb in LLC lung tumor versus healthy lung cDCs1, but not in splenic cDCs1, suggested that cDCs1 undergo lysosomal activation and stress in tumours (**Fig. 1d, 1e**). By confocal microscopy we identified Gpnmb on the membranes of tumour cDC1 lysosomes (**Fig. 1f**). LysoTracker probes label lysosomes, as well as late endosomes and for this reason they cannot be safely used as a measure of lysosome pH ^45^. The fluorescent probe DQ-BSA is internalized by endocytosis and degraded by proteolysis only in lysosomes, emitting fluorescence upon degradation ^45^. Compared to purified healthy lung cDCs1, lung tumour cDCs1 consistently showed a higher DQ-BSA fluorescence signal at various time points, suggesting that the lysosomes of tumour cDCs1 had a higher degradation capacity (**Fig. 1g**). cDCs1 must slowly digest dead cancer cell debris for efficient cross-presentation ^10,29^. To monitor the cancer antigen degradation capacity of tumor cDC1s, we FACS-sorted tumour and healthy lung cDCs1 and co-cultured them with tagged apoptotic LLC cells. The antigen load of tumour and healthy cDCs1 peaked on the same day but was consistently lower in tumor cDC1s at different time points (**Fig. 1h**). This was rather due to faster antigen degradation by tumour cDCs1 rather than lower endocytosis, as an endocytosis fluorescent reporter (FITC-Dextran, 40.000 MW) showed no difference between lung tumour and healthy lung cDCs1 (**Fig. 1i**).

The nutrient sensor mTOR forms the catalytic core of the mTORC1/2 complexes and controls lysosomes ^46^(**Fig. 1j**). In nutrient and growth factor rich environments mTOR phosphorylation downstream of AKT activates mTORC1 to enhance protein synthesis and suppress lysosomal nutrient degradation, by sequestering the master regulator of lysosome biogenesis TFEB outside the nucleus ^47^. Upon starvation, dissociation of mTOR from mTORC1 allows TFEB translocation to the nucleus, lysosome biogenesis and enhanced catabolic processes ^47^. FACS analysis gating on cDCs1 showed lower levels of phospho-mTOR in lung tumor versus healthy lungs cDCs1, while phospho-AKT did not differ between the two (**Fig.1j**), suggesting that the signaling pathway that drives mTOR inactivation in tumour cDC1 is AKT-independent. To investigate the specificity of tumour-induced lysosomal states for cDC1, we assessed lysotracker stain and intracelluar Gpnmb in tumour and healthy lung cDCs2 and macrophages. Endo-lysosomal acidification was induced at low levels in tumour cDC2, but not in macrophages, while only cDCs1 showed lysosomal stress, measured by Gpnmb (**Suppl. Fig. 5**). To directly show that the tumor microenvironment activates DC1 lysosomes, we exposed the MutuDC1 cell line to Tumor Culture Medium (TCM) versus Healthy Lung Culture Medium (HCM). Upon TCM exposure, MutuDC1 acquired a lysosomal function profile that was similar to that of tumour cDC1 and was suppressed by the mTOR activator bafilomycin A1 (**Suppl. Fig. 6**). Phosphoproteomics analysis of TCM-exposed versus HCM-exposed MuTucDC1 failed to identify an inhibitory signaling pathway upstream of mTOR (**Suppl. Fig. 7**). Collectively, building on transcriptomics for unbiased hypothesis generation we discovered a functional module which points to lysosomal processes as key determinants of tumour-induced cDC1 states.

### Lysosomal genes of cDCs1 are transcriptionally upregulated by type I IFNs

To infer the regulons that drive lysosomal states in tumors we applied the scalable SCENIC platform in the murine scRNAseq dataset ^48^. The activity of the predicted regulons in all individual cDCs1 was quantified and cellular AUCs were used as input for UMAP visualization and clustering. The SCENIC AUC UMAP clearly separated a tumor-specific cDC1 cluster that almost completely overlapped with the C1 tumor-specific cluster in the transcriptome UMAP **(Fig. 2a)**. Ranking all regulons of the tumor C1 cluster according to the regulon specificity score pointed to *STAT1, STAT2 and IRF7* as the top regulons **(Fig. 2a)**. These transcription factors are major activators of transcription of interferon-stimulated genes. Type I IFNs drive cDC1 maturation and migration ^21,49,50^ and metabolic processes are critical determinants of these functional properties of cDCs ^51^. Type I IFNs regulate mTOR ^52,53^ and cDCs use mTOR to spatio-temporally adapt their metabolism to the diverse metabolic needs of antigen processing, maturation and migration to dLNs ^54,55^. However, the interplay between type I IFNs, mTOR and lysosomes is unclear. First, we performed in vitro assays with the MutuDC1 line. Amino acid deprivation of MutuDC1 or exposure to pure type I IFNs, but not to IFNγ, increased intracellular Gpnmb, endo-lysosome acidification and lysosome biogenesis **(Suppl. Fig. 8)**. IFNβ-induced lysosomal responses were suppressed by the mTOR activator Bafilomycin A1 **(Suppl. Fig. 8)**. To validate these findings in vivo, we generated mixed bone marrow chimeras by sub-lethal irradiation and transplantation of CD45.1 wild type and CD45.2 INFRA1 knockout (KO) bone marrow cells at a 1:1 ratio **(Fig. 2b)**. Chimeric mice received lung injection of LLC cells. Tumors were excised, digested and intratumoural cDCs1 were analyzed by FACS, gating on CD45.1 wild type (WT) and CD45.2 INFRA1^KO^ cDC1s. INFRA1^KO^ intratumoral cDCs1 showed decreased endo-lysosome acidification compared to WT cDC1s and increased phospho-mTOR **(Fig. 2b)**. RNAseq analysis of FACS-sorted intratumoural cDCs1 showed a higher number of downregulated versus upregulated genes in IFNAR1 KO cDCs1 **(Fig. 2c)**. Among top downregulated were genes involved in IFN responses (*MyD88, Irf7, Oas1a, Ifi27*) and cDC1 maturation/migration (*CCR7, IL12b*), as expected ^21,49,50^ **(Fig. 2c)**. Critically, the lysosomal enzymes *Neu1, Lgmn, Ctsa* and *Ctse* and the Golgi-to-lysosome trafficking mediator *Gga2*, were also found highly down-regulated (**Fig. 2c**), suggesting a positive effect of type I IFNs in lysosomal activities at the transcription level. Likewise, down-regulated genes in IFNAR1 KO cDCs1 showed significant enrichment in metabolic/lysosomal processes, viral responses and cell migration (**Fig. 2d**). To reconstruct the gene regulatory networks that control IFNAR-driven cDC1 states we performed gene regulatory network inference using GENIE3 and all 195 significantly down-regulated genes as input ^56^. Similar to what we had discovered in the scRNAseq dataset using the SCENIC algorithm, GENIE3-predicted network topology revealed a lysosome-associated module that was co-regulated by several TFs, among which the Interferon Regulatory Factor *IRF7* (**Fig. 2e**). To further corroborate that IFN-induced TFs regulate transcription of lysosomal genes, we curated a list of lysosomal processes-related genes **(Suppl. Table 1)** and performed TF enrichment analysis (Chea3) and TF binding motif analysis (RcisTarget). STAT1 and IRF7 binding motifs were over-represented in our lysosomal gene list and **(Fig. 2f),** while STAT1, STAT2 and IRF7 were among top-ranked upstream TFs **(Suppl. Fig. 9)**. In conclusion, by applying cytokine exposure assays, in vivo cytokine receptor perturbations and bioinformatics tools, we discovered that type I IFNs lead to transcriptional activation of cDC1 lysosomal genes in tumours.

**Figure 2.**
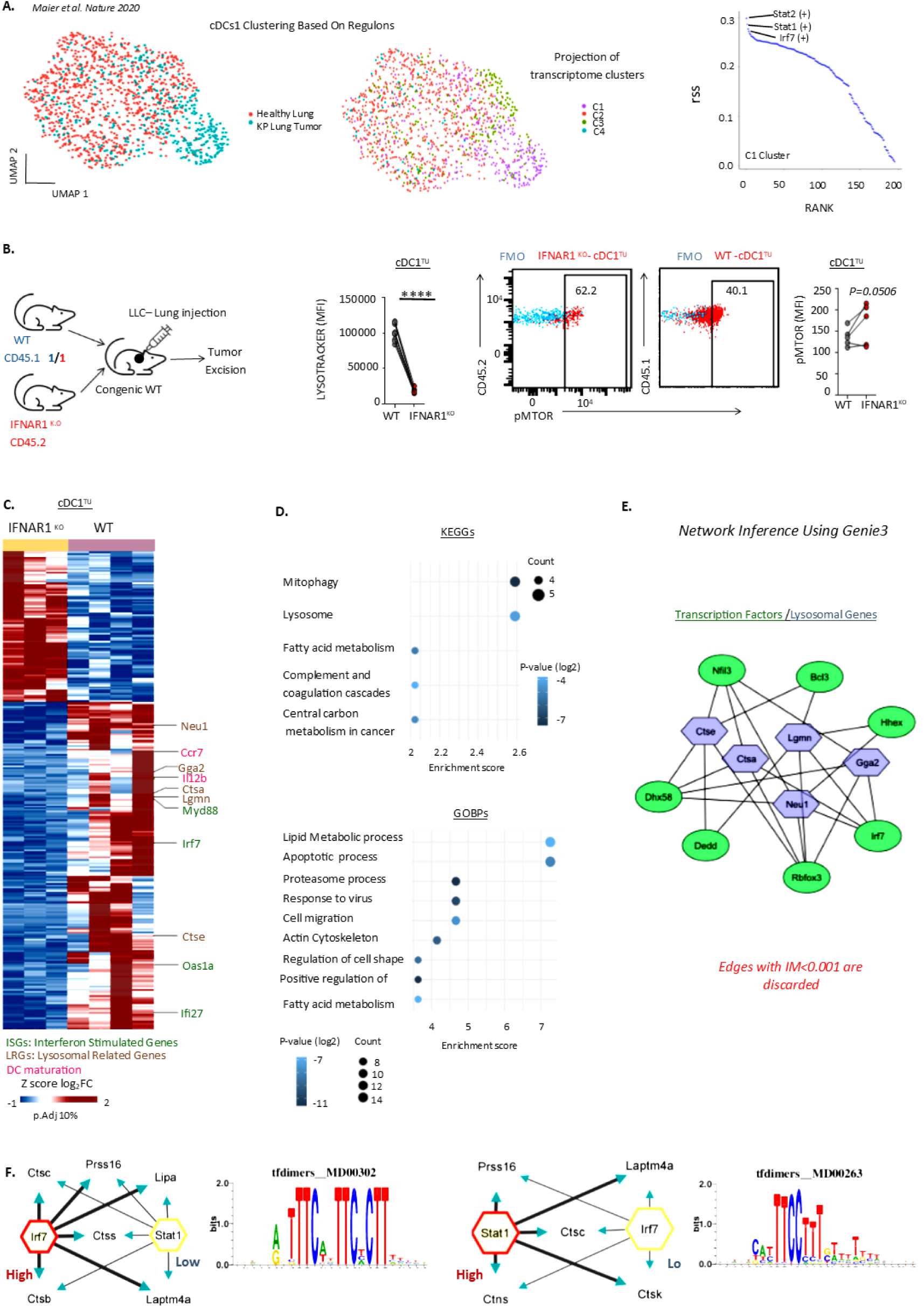
Type I IFNs regulate cDC1 lysosomes at the transcription level. **A.** Regulatory network inference and clustering of KP lung tumor and healthy lung cDCs1 using SCENIC ^48^. Left to right. The activity of the predicted regulons in individual cDC1s was quantified and cellular AUCs were used as input for UMAP. UMAP color-coded based on tissue of origin or cDC1 transcriptome clusters. Graph depicting ranking of the regulon specificity scores of the tumor-specific cluster C1. **B.** LLC lung tumours of mixed bone marrow chimeric mice (CD45.1 wild type / CD45.2 IFNAR1^KO^) were excised and digested. Lysotraker and phospho-mTOR were analyzed by FACS gating on IFNAR1^KO^-cDC1^TU^ and WT-cDC1^TU^. Error bars, mean± sem; **** p<0.0001, two-tailed paired t-test. MFI: Mean Fluorescence Intensity. **C.** IFNAR1^KO^-cDC1^TU^ and WT-cDC1^TU^ (n=4 biological replicates per condition) were analyzed by bulk-RNAseq. Heatmap of 285 differentially expressed genes. **D.** KEGG pathways and Gene Ontology (GO) enrichment analyses of 195 down-regulated genes in IFNAR1^KO^-cDC1^TU^. **E.** Cytoscape view of network inference using GENIE3 ^56^. Analysis of top 195 predicted interactions (IM >0.001) revealed a module that corresponds to lysosomal genes. **F.** Top high confidence enriched motifs for STAT1 and IRF7 in the regulatory regions of selected lysosomal genes (RcisTarget). High: High confidence, Low: Low confidence.

### Type I IFN-induced permeabilization of lysosomal membranes causes cDC1 death and constricts anti-tumour immunity

Lysosomal gene transcription can be induced by lysosomal membrane permeabilization (LMP) as a balancing mechanism to generate new lysosomes ^31,46^. Failure to repair LMP or remove the damaged lysosomes results in the uncontrolled release of lysosomal enzymes and induces a type of death known as lysosome-dependent cell death (LDCD) ^31^. We hypothesized that LDCD might explain cDC1 extinction from tumor tissues and that lysosomal activation of cDCs1 in tumours may be due to LMP. First, we quantified cDCs1 in our orthotopic lung cancer model. We excised primary murine LLC lung tumors and healthy lungs, stained single cell suspensions with antibodies against cDC markers and performed FACS. We concatenated lineage negative, MHCII^+^CD11c^+^ cDCs of tumor and healthy samples and performed dimensionality reduction and unbiased clustering. cDCs separated in 4 clusters: an XCR1^high^CD11b^-^bona fide cDC1s, XCR1^-^CD11b^+^ bona fide cDC2 and two intermediate states XCR1^low^CDC11b^-^ (XCR1low mReg cDC) and XCR1^-^ CD11b^-^ (XCR1-mReg cDC) (**Fig. 3a**). Bona fide tumor infiltrating cDCs1 were less in lung tumors versus healthy lungs (**Fig. 3a**). In accordance, cDC1 density was decreased in KP lung tumours (**Fig. 3b**). Quantification of dead cDCs1 in tissues was not feasible, because antibodies bind non-specifically to dead cells and dead cDCs1 cannot be discerned safely from other dead cells. Exposure of phosphatidylserine at the cell surface, detected by annexin V stain, is an early feature of cells undergoing apoptotic and non-apoptotic cell death^57^. Quantification of Annexin V positive cDCs1 by FACS showed a higher percentage of cDCs1 in lung tumors versus healthy lungs **(Fig. 3c)**. In addition, approximately 50% of cDCs1 that were found alive in lung tumors of wild type mice were IFNAR1 positive, while live NFRA1 positive cDCs1 infiltrating healthy lungs were around 80%, further supporting our hypothesis that type I IFNs eliminate cDC1 **(Fig. 3d)**. Next, we exposed the MutuDC1 line to lung Tumor Culture Medium (TCM), Healthy lung Culture Medium (HCM), or pure IFNs and again analyzed Annexin V/Dead staining. A lower percentage of TCM and IFNβ exposed-MutuDC1 versus HCM and IFNγ-exposed MutuDC1, respectively, were alive **(Suppl. Fig. 10)**. Cytosolic release of lysosomal enzymes produces a diffuse fluorescence pattern that is specific for LMP. Cathepsin D was imaged outside the lysosomes of TCM-exposed MutuDC1, suggesting that they underwent LMP **(Suppl. Fig. 10)** ^31^. Galectin 3 translocation at the membrane of leaky lysosomes creates a puncta stain that is highly sensitive and specific for LMP ^31,45,58^. Lung tumour cDCs1, as well as TCM-exposed and IFNb exposed MutuDC1, showed extensive galectin 3 puncta in almost all cells, suggesting severe LMP **(Fig. 3e)**. To show in vivo that type I IFNs induce cDC1 LMP in tumors, we analyzed lung tumours and spleens of CD45.1 WT/CD45.2 IFNAR1^KO^ chimeric mice. INFRA1^KO^ cDCs1 outnumbered WT cDC1s in tumors, but not in the spleen, while Annexin V stain was decreased, paralleled by an impressively decreased Galectin 3 puncta **(Fig. 3f)**. LMP can trigger other cell death signaling pathways, such as apoptosis or pyroptosis ^31^. Lysosomal protease inhibitors are used to determine whether LMP is the primary or secondary cause of cell death ^31^. Accordingly, MutuDC1 was rescued by IFNβ-induced death by the protease inhibitor leupeptin **(Fig. 3g)**. To replicate this effect in vivo, we co-administered pooled leupeptin-treated and PBS-treated MutuDC1 to flank tumour bearing mice via direct intratumoural injection, excised tumours and analyzed MutuDC1 by FACS. Leupeptin pre-treatment decreased MutuDC1 lysosomal stress, measured by Gpnmb intracellular stain, accompanied by a relative increase in their numbers **(Fig. 3h)**. Importantly, when leupeptin-treated versus PBS-treated MutuDC1 were administered in array, flank tumours of the first grew at slower rates (**Fig. 3i**). Collectively, in vivo and ex vivo orthogonal experimental approaches provide solid evidence that type I IFNs induce LDCD in cDCs1 in tumours and point to cDC1 LMP as a novel immunotherapeutic target

**Figure 3.**
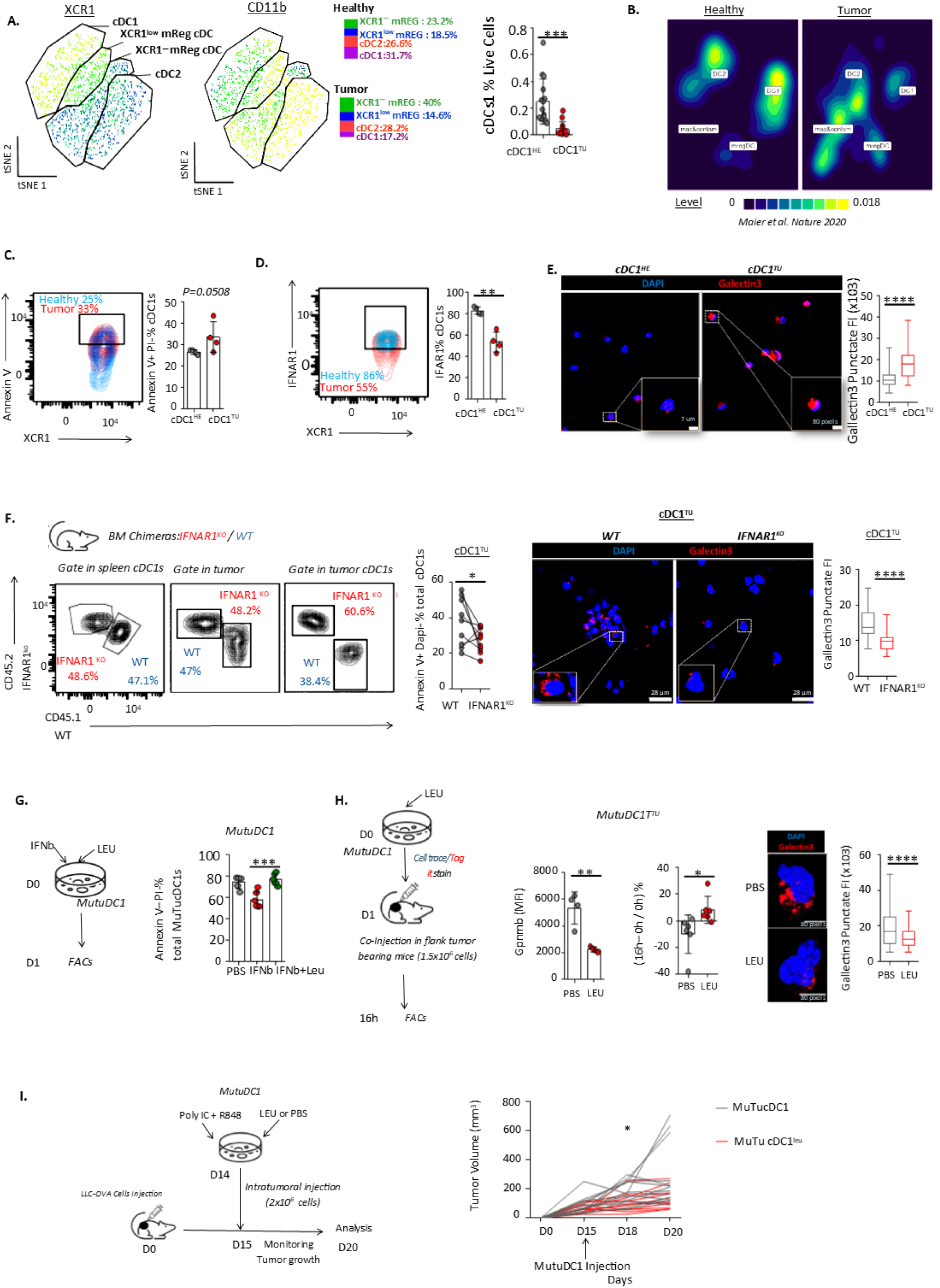
Type I IFNs induce lysosomal membrane permeabilization (LMP) and death, limiting anti-tumour immunity. **A-F**. LLC cells was injected on the left lung lobe of mice. Lung tumors and healthy lungs of control mice were excised, digested and analyzed by FACS. **A.** Left to right. Lin^-^CD11c^+^MHCII^+^ cDCs from different samples were concatenated into one FCS file and t-SNE was run using XCR1 and CD11b as input. cDC1 frequency in lung tumours (cDC1^TU^) versus lungs (cDC1^HE^). **B.** Density plot of cDC subsets in a murine lung/KP lung tumour scRNAseq dataset ^42^. **C.** Percent annexin V^+^ live cDCs1 quantified by FACS. **D.** INFAR1 percent cDC1. **E.** cDC1^TU^ and cDC1^HE^ were FACS sorted and Galectin 3 puncta was quantified by image analysis (n=89 cells per condition). **F.** LLC cells were injected in the left lung lobe of mixed bone marrow chimeras (CD45.1^WT^ / CD45.2 INFRA1^KO^). Spleens and lung tumors were excised and digested. Left to right. FACS plots depict WT to IFNAR1^KO^ among total CD45^+^ cells or cDCs1. Percent annexin V^+^ live WT-cDC1^TU^ versus IFNAR1^KO^-cDC1^TU^. Galectin 3 puncta of FACS-sorted WT-cDC1^TU^ versus IFNAR1^KO^-cDC1^TU^ quantified by image analysis (n=33 cells per condition). **G.** MutuDC1 was exposed to IFNβ in the presence or absence of the protease inhibitor leupeptin. Percent annexin V^-^ live MutuDC1 quantified by FACS. **H.** From left to right. Leupeptin-treated MutuDC1^leup^ and PBS-treated MutuDC1^pbs^ were co- injected in LLC flank tumours (0h). 16h later tumours were excised and digested. Percent increase in intratumoural MutuDC1 subsets enumerated by FACS. Intracellular Gpnmb at 16h. Galectin 3 puncta in intratumoural FACS-sorted MutuDC1^leup^ and MutuDC1^pbs^ quantified by image analysis (n=107 cells per condition). **I.** Leupeptin-treated MutuDC1^leup^ versus PBS-treated MutuDC1^pbs^ were injected in LLC flank tumours. Tumour volumes were measured every other day. On d7 tumours were excised, digested and analyzed by FACS. **A-I**. Representative or cumulative data from at least 2 independent experiments with 4-6 biological replicates per group per experiment. Each biological replicate of FACS- sorted cDCs1 was pooled from 3-5 mice. *p<0.05, **p<0.01, ***<0.001, ****<0.0001, two-tailed unpaired or paired t-test. Error bars, mean± sem.

### Murine tumour cDC1 lysosome states can be replicated across four different human cancer types

Humans are much more diverse and heterogeneous compared to laboratory mice. To investigate the translatability of our murine studies in the human system we leveraged a publicly available mega-integrated dataset consisting of 178,651 human mononuclear phagocytes across 41 datasets ^59^. We filtered out non-cancer patients and patients with malignancies of lymphoid tissues, leaving us with 2315 annotated cDCs1 across 39 paired human and cancer samples of the liver, lung, colon and kidney. cDC1 density decreased in tumour tissues **(Suppl. Fig. 11a)**. Differential expression analysis showed a small number of deregulated genes in tumour states. Among the few genes that were up-regulated were IFN-inducible genes (STAT1, GBP2) and cathepsins (CTSA, CTSD) **(Suppl. Fig. 11b)**. Accordingly, pathway analysis showed significant enrichment in GOBP and KEGG pathways related to IFN responses, lysosomal processes and apoptosis **(Suppl. Fig. 11c)**. Regulon inference using the SCENIC algorithm pointed to STAT1 as the most important transcription factor regulating the gene expression profile of tumor cDCs1 **(Suppl. Fig. 11d)**. Furthermore, purified IFNβ directly enhanced the endo-lysosome acidification of cDCs1 of healthy donors **(Suppl. Fig. 10e)**. Thus, transcriptomics and functional profiles of human cDCs1 suggest that similar to their murine counterparts, they are likely to undergo lysosomal stress in tumours, driven by type I IFNs.

### IFN-inducible GBP-2 induces LMP and can be targeted to boost DC1 therapy

LMP is known to be induced by lysosomal stress in the presence of pore-forming molecules, such as Bax and Bac ^31,46^. GBPs are IFN-inducible GTPases that can form supramolecular complexes on the membrane of pathogen-containing vacuoles and on the bacterial membrane, leading to vacuolar/bacterial lysis ^9,60^. Interestingly, GBP-2 was one of the most highly upregulated genes in human and murine tumour cDC1 transcriptomics datasets (**Fig. 4a**). Membranes of healthy lysosomes are normally protected from GBPs ^9^. We hypothesized that GBP-2 overexpression driven by type I IFNs in the tumour microenvironment might induce GBP-2 accumulation at the membrane of stressed lysosomes and LMP. We stained IFNβ versus PBS-exposed MutuDC1 with antibodies against GBP1-5 and LAMP-3, a lysosome-associated membrane protein. Confocal imaging suggested that IFNβ drove strong co-localization of GBPs with LAMP-3 on MutuDC1 lysosomes, which was validated by electron microscopy (**Fig. 4b, 4c**). Next, we infected MutuDC1 stably expressing dCAS9-KRAB with BFP-expressing lentiviruses carrying either of two sgRNAs targeting GBP-2 (or sgRNA control), assessed the level of GBP-2 repression and selected the most efficiently repressed MutuDC1 for expansion. Consistent with our initial hypothesis, GBP-2 repression increased viability of MutuDC1 in culture and further protected MutuDC1 from IFNβ-driven death (**Fig. 4d**). Importantly, IFNβ-exposed sgGBP2-dCAS9-KRAB MutuDC1 showed drastically decreased Galectin 3 puncta compared to sgCTL-dCAS9-KRAB MutuDC1 (**Fig. 4e**). Thus, type I IFNs induce LMP in MutuDC1 via GBP-2 in vitro. To validate our hypothesis in vivo, we co-administered pooled sgGBP2-dCAS9-KRAB MutuDC1 and sgCTL-dCAS9-KRAB MutuDC1 to flank tumour bearing mice via direct intratumoural injection, excised tumours and analyzed MutuDC1 numbers by FACS. GBP-2 repressed MutuDC1 outnumbered controlled MutuDC1 in flank tumours **(Fig. 4f)**. We hypothesized that boosting MutuDC1 viability in tumours might enhance its anti-tumour efficacy and help overcome resistance to immunotherapy. We designed a combination cell therapy approach where sgGBP2-dCAS9-KRAB MutuDC1 and OVA-specific CD8 T cells were co-injected intratumourally in OVA-LLC flank tumour bearing mice (**Fig. 4g**). In parallel, mice were administered the checkpoint inhibitor aPDL1. Flank tumours of sgGBP2-dCAS9-KRAB MutuDC1 versus sgCTL-dCAS9-KRAB MutuDC1 recipients grew at slower rates and were infiltrated by increased cancer antigen-specific T cells, shown by PD1, ICOS and H-2Kb OVA tetramer staining (**Fig. 4g**). Therefore, type I IFN-induced LMP is driven by GBP-2 and GBP-2 repression boosts the efficacy of DC1 therapies.

**Figure 4.**
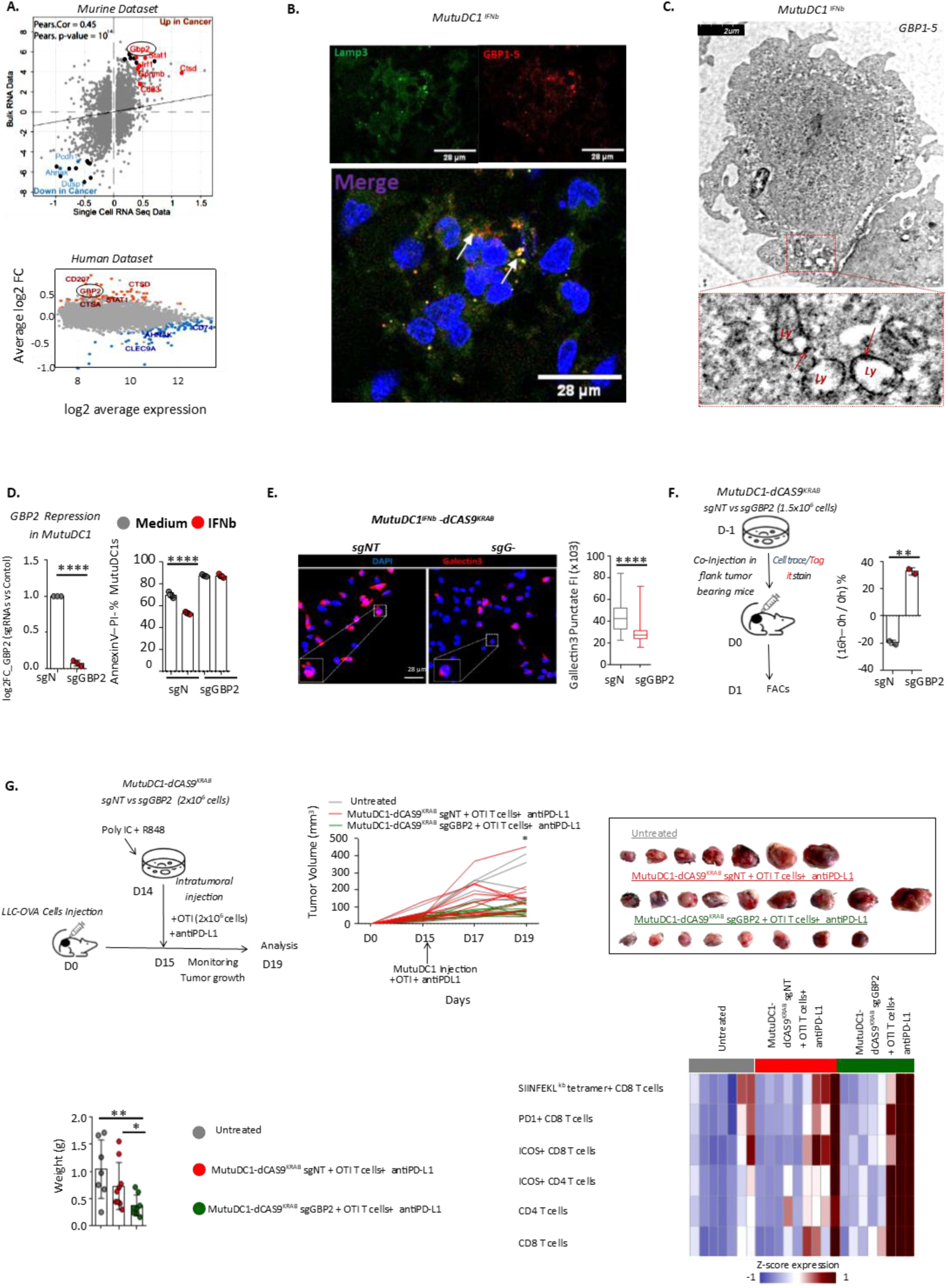
Type I IFN-induced LMP is driven by GBP-2 and targeting GBP-2 improves the efficacy of DC1 therapy. **A**. Up. Correlation analysis of log2 fold change values obtained from murine bulk RNA-Seq (in-house dataset) and scRNAseq (public dataset^42^) for all genes in the comparisons of murine lung tumour cDC1 versus healthy lung cDC1. Down. Mean-Average(MA)-plot of log2 average expression versus log2 fold change between human tumour cDC1 versus paired unaffected tissue cDC1 (public dataset) ^59^. Colored dots indicate genes that are significantly differentially expressed. **B.** Co- localization of GBP1-5 and lysosomal membrane protein LAMP-3 was analysed in Tumour Culture Medium-treated MutuDC1^TCM^ . **C.** Immunoelectron microscopic localization of GBP1-5 in IFNβ-treated MutuDC1. GOLD labelling is noted in the membrane of lysosomes (Ly). **D.** Percent annexin V^-^ live in IFNβ/PBS-treated sgGBP2- dCAS9^KRAB^ MutuDC1 and sgNT-dCAS9^KRAB^ MutuDC1. Pooled data from 2 independent experiments. **E.** Galectin 3 puncta in FNβ-treated sgGBP2-dCAS9^KRAB^ MutuDC1 versus sgCTL-dCAS9^KRAB^ MutuDC1 quantified by confocal imaging (n=90 cells per condition). **F.** sgGBP2-dCAS9^KRAB^ MutuDC1 versus sgCTL-dCAS9^KRAB^ MutuDC1 were co-injected in LLC flank tumours (0h). 16h later tumours were excised and digested. Percent increase in intratumoural MutuDC1 subsets enumerated by FACS. Data are representative of 2 independent experiments. **G.** sgGBP2-dCAS9^KRAB^ MutuDC1 versus sgCTL- dCAS9^KRAB^ MutuDC1 were injected in LLC flank tumours. Tumour volumes were measured every other day. On d4 tumours were excised, digested and analyzed by FACS. Representative data from 2 independent experiments, n=7-10 mice per group, are shown. Numbers of intratumoral T cells were normalized to numbers of intratumoral cancer cells. **D-G**. *p<0.05, **p<0.01, ****<0.0001, two-tailed unpaired or paired t-test or ANOVA. Error bars, mean± sem.

## Discussion

cDC1s are scarce in solid tumors, which facilitates immune escape, confers immunotherapy resistance and decreases patient survival ^4–7,18,19,23–26^. Here we found that cDC1 tumour deserts are due to type I IFN-induced LDCD. The cascade of events that compromises cDC1 survival stems from aberrant transcription of a set of genes that belong to the endo-lysosomal damage response pathway ^31^ and of an IFN-inducible GTPase, i.e. GBP-2. GBP-2 accumulated in the membrane of the cDC1 lysosomes and induced LMP, allowing the release of the amplified protease pool to the cytosol and death. We speculate that GBP-2 is part of the pore-forming complex that allows translocation of the antigens from the phago-lysosome to the cytosol for cross-presentation. This physiological process might well-controlled in steady state cDCs1 due to their intrinsically low proteolytic activity ^10,29^ but it turns out to be lethal upon IFN I-driven lysosomal destabilization and GBP-2 overexpression. Targeting LDCD by neutralizing the proteolytic activity of proteases or repressing GBP-2 increased the therapeutic effects of adoptive cDC1 transfer pointing to LDCD as a novel immunotherapeutic target.

Lysosomal activation can be induced by starvation, oxidative stress or as part of the lysosome stress response pathway to induce biogenesis of new healthy lysosomes ^31,46^. The latter is induced by lysosomal dysfunction, such as deficiency in lysosomal enzymes in lysosomal storage disorders or lysosomal membrane permeabilization (LMP) ^31,46^. LMP can be triggered by endocytosed material (e.g. urate crystals or myelin debris) or bacteria that rupture phagolysosome membranes to escape into the cytosol ^31,61^. LDCD results from inability of the endo-lysosomal damage response pathway to repair LMP, allowing the excessive release of cathepsins into the cytosol ^31,46^. Minor damage to the membrane is repaired by the Endosomal Sorting Complexes Required for Transport (ESCRT) machinery and by oxysterol-binding protein (OSBP)-related proteins (ORPs) ^58,62^. Higher damage initiates the clearance of terminally damaged lysosomes by autophagy, termed lysophagy ^31,46,61^. The above pathways are largely coordinated by galectin 3 ^63^. In parallel, galectin 8, also recruited to the damaged lysosome membrane, dissociates the mammalian Target of Rapamycin (mTOR) leading to dephosphorylation of transcription factor EB (TFEB) and its translocation to the nucleus, where it induces transcription of lysosomal genes for the biogenesis of new lysosomes ^64^. Two lines of evidence indicate that primary cDCs1 undergo IFN I-driven LMP in tumours and that they have active endo-lysosomal damage response pathway: i) galectin 3 puncta on damaged lysosomes were observed in IFNAR1 competent but not IFNAR1 KO murine intratumoural cDCs1 and, ii) cathepsins were identified outside the lysosomes of TCM-exposed MutuDC1. We also directly showed that tumour culture media induce LMP and LDCD via type I IFNs in vitro. LMP can trigger other cell death signaling pathways, such as apoptosis or pyroptosis ^31^. Targeting LMP by the lysosomal protease inhibitor leupeptin or by GBP-2 repression, increased cDC1 viability, confirming LDCD as the primary type of cDC1 death in tumours^13,31^.

Type I IFNs can induce lysosome activation, rupture of the pathogen-containing vacuole and cell death in Salmonella infected intestinal epithelial cells ^65^. We discovered that LMP/LDCD in cDCs1 were paralleled by increased lysosome acidification and degradative capacity and decreased phospho-mTOR. It could be that an unknown mediator of the IFNAR1 signaling pathway directly inhibits mTOR or activates mTOR inhibitors, albeit we could not deduce such a mechanism from our phosphoproteomics analysis. We did, however, discover binding motifs for IFN-inducible TFs in promoters/enhancers of lysosomal genes. Furthermore, IFNAR1 KO-WT mixed bone marrow chimeras showed that type I IFNs up-regulate TFs with binding specificities for regulatory regions of lysosomal genes. Thus, there seems to be a direct transcriptional link between type I IFNs and lysosome activation. We also suspect a positive feedback loop, whereby increased lysosomal proteolytic activities aggravate LMP toxicity, which in turn further activates lysosomes via the galectin 8/mTOR axis, worsening LMP ^64^. Indeed, type I IFN-induced activation was decreased by the mTOR activator Bafilomycin in vitro. Conversely, we have observed that mice that lack the mTOR inhibitor PTEN show increased intratumoural cDCs1 ^42^. mTOR is primarily a nutrient sensor inhibited by starvation, as we confirmed in vitro for cDCs1 ^46^. Thus, the functional module that consists of IFN-inducible TFs, transcriptional activation of lysosomes and mTOR suppression, depends on type I IFNs, but is likely further amplified by the endo-lysosomal damage response pathway and tumour-associated starvation.

Self-membranes are thought to be either protected from GBPs by GBP inhibitors (e.g. IRGMs) or GBPs recognize an unknown molecule on the surface of pathogen-containing vacuoles that does not exist on self-membranes ^9^. We discovered that under lysosomal stress, GBPs accumulated in the membrane of the cDCs1 lysosomes, causing LMP and LDCD. Considering that self-membrane bound GBPs have been very rarely observed ^9^, it is puzzling why lysosomal membranes of cDCs1 are GBP targets. cDCs1 are the most potent cross-presenting cells and the only DCs that can release antigens to the cytosol after phagosome-lysosome fusion via an unknown pore complex ^10,30^. We believe that GBP-2 is key constituent of this unknown complex. Thus, IFN I-driven lysosomal destabilization and GBP-2 overexpression may render a physiological mechanism of antigen translocation for cross-presentation to a death pathway.

A recent study reported a progressive decrease in cDCs1 in lung tumor draining LNs ^36^. Both lysosomal stress ^66,67^ and increased GBP2 ^68^ that we have discovered can change cytoskeleton dynamics and calcium signaling in cDCs1, decreasing their migration ^69^. Accordingly, among the top down-regulated genes in our tumor cDC1 dataset there were several related to cytoskeleton and cell adhesion. Overall, impaired cDC1 migratory capacity, coupled by a progressive decrease in numbers of intratumoural cDCs1 due to LDCD, may account for a progressive decrease in nodal cDCs1.

Only two studies have targeted cDC1 IFN signaling in vivo in cancer ^21,50^. Type I IFNs and IFNγ activate similar downstream signaling cascades, critically mediated by the TF IRF1. Conditional IRF1 deletion impairs cDC1 maturation and chemo-cytokine expression, increasing tumour growth ^21^. However, targeting IFNAR1 -by contrast to IFNγR-in cDCs1 has no impact on tumour growth, suggesting that the beneficial effects of IFNs on cDC1-driven anti-tumour immunity are mediated by IFNγ ^50^. In our IFNAR1 KO/WT mixed bone marrow chimeras, IFNAR1 KO cDCs1 were protected from LDCD and increased in numbers in tumours. These seemingly contradictory data indicate that type 1 IFNs are a double edge sword, increasing cDC1 maturation and cross-presentation on the one hand ^21,49,50^, decreasing their viability due to LDCD on the other.

Although type I IFNs have been recently implicated in immunotherapy resistance and T cell exhaustion ^70,71^, they are generally recognized as powerful immunostimulating anti-tumour agents ^49,72–74^. Thus, it is difficult to envisage a therapeutic scenario whereby targeting IFNs per se to prevent cDC1 LMP will ameliorate clinical outcomes of cancer patients. There are already research efforts to directly target LMP to treat lysosomal storage disorders ^31^. Such drugs could be also re-purposed to prevent cDC1 LMP in tumors. Our pre-clinical experiments with the administration of MutuDC1 pre-treated with the protease inhibitor leupeptin or GBP-2 deficient MutuDC1 were very promising. In summary, cDCs1 are fundamental for cancer immunity, but due to their scarcity they have not been harnessed for therapy to date. We have discovered that they undergo LMP and LDCD in tumours, which causes or contributes to their extinction, with detrimental outcomes for anti-tumour immunity. cDC1 LMP/LDCD is treatable via protease inhibition and GBP-2 repression and we expect more molecular targets to be uncovered as we explore more the mechanisms of this discovery.

## AKNOWLEDGEMENTS

We acknowledge G. Kollias, E. Andreakos, and G. Stathopoulos for sharing mouse models and cell lines. The NIH Tetramer Core Facility for SIINFEKL tetramers. We acknowledge V.Zarkou, M. Papanastasatou and Papadogiorgaki Sevasti for sharing technical expertise. We thank BSRC “Al. Fleming” flow cytometry, animal house, genomics facilities and the single cell analysis unit. We thank Mosaiques diagnostics for mass spectrometry. We thank EMBL GeneCore for library preparation and Next Generation Sequencing. This project has received funding from the H2020 research and innovation programme (ERA PerMed JTC2019), the General Secretariat of Research and Innovation (Τ11ΕΡΑ4-00044, BronchoBOC, MIS 5070471) and by NextGenerationEU, under the call 16971 RESEARCH -CREATE - INNOVATE (TAEDK-06185). E. Aerakis is supported by the Hellenic Foundation for Research and Innovation (HFRI) under the 3rd Call for HFRI PhD Fellowships (No 6141). I. Angelidis is supported by the Europeans Respiratory Society and the EU’s H2020 research and innovation program under a RESPIRE4 Marie Sklodowska-Curie Actions grant agreement (No 847462).

## CONTRIBUTIONS

Conception, study design, M.T.; Development of methodology, E.A., M.T.; Acquisition of data, E.A., A.C., M.A., M.M.P., I.A., D.Koum., A.G, P.H., D.Ker., M.Mak., A.V., B.M., S.H., M.Mer., M.T.; Analysis and interpretation of data, E.A., B.M., S.H., M.Mer., M.T.; Writing the manuscript, E.A., M.T.; Preparing the figures, E.A., A.C.; Study supervision, M.T.

## METHODS

### Cell lines

The Lewis Lung Carcinoma cell line (LLC), obtained from American Type Collection Cultures (Manassas, VA) and the C57BL/6-derived urethane-induced lung adenocarcinoma CULA cell line, provided by G. Stathopoulos (Helmholtz Zentrum München), transduced with zsGreen and/or OVAmCherry and/or Luc-ZsGreen lentiviruses and sorted based on fluorescent protein expression, have been previously described ^1,2^. The MutuDC1940 cell line developed by Acha-Orbea Hans ^3^ was provided by Caetano Reis E Sousa. Cancer cells were maintained in DMEM, containing 10% heat-inactivated FBS, 1% L-glutamine, and 1% penicillin/streptomycin (Gibco). MutuDC1 was maintained in IMDM-GlutaMAX, 10% heat-inactivated FBS, 1% penicillin/streptomycin, 25 mM HEPES, 3.074 g/L Sodium Bicarbonate and 50 µM β-mercaptoethanol. All cell lines were regularly tested negative for the presence of mycoplasma contamination using a PCR-based technology.

### sgRNA design and preparation

The sgRNAs were designed using the Benchling platform. Total length of sgRNAs was 20nt, the first nucleotide was G and is designed for CAS9 that recognize NGG PAM site. The target site that recognizes sgRNAs was located from -50bp to +100bp between TSS. To prepare each sgRNA, the sgRNA (IDT) was reconstituted to 100 μM ddH_2_O. To prepare the sgRNA duplexes, the two oligonucleotides were mixed at equal concentrations (100 μΜ) in a sterile PCR tube containing annealing buffer (10mM Tris pH 8.0, 50mM NaCl and 1mM EDTA pH 8.0). Oligos were annealed by heating at 95°C for 5 min followed directly by hybridization for 15 min at room temperature in a PCR thermocycler. The mix was then placed on ice or frozen at −20°C until further use. sgRNAs were cloned into the pLG1-puro non-targeting sgRNA 1 backbone between BstXI and BlpI restriction sites. The following sgRNAs were used: GBP2 forward (Fw): 5’- CATAGACCCTGTAGTAACCC -3’ and reverse (Rv): 5’- GGGTTACTACAGGGTCTATG -3’

### Transfection and transduction

For lentiviral production, HEK lenti-X 293 T cells were seeded in a 10-cm dish. Upon reaching 70% confluence, they were transfected with a mixture of 16μl JetPRIME (Polyplus), 2.6 μg psPAX2 (addgene #12260), 1.4 μg pMD2G (addgene #12259) and 3.75 μg pLV plasmid coding for the protein of interest (pHR-SFFV-KRAB-dCas9-P2A-mCherry, addgene #60954) or (pLG1-puro Non-targeting sgRNA 1, addgene #109002) in 500μl JetPRIME buffer. 6 h after transfection the transfection mixture –containing culture medium was replaced with fresh one. On day 3 after transfection, the pseudotyped virus-containing culture medium was collected and filtered through 0.44-μm filter. Aliquots of virus were stored at -80°C

For lentiviral transduction of MutuDC1 0.4 x10^6^ cells were seeded in six well plate. Next the pseudotyped virus-containing culture medium was diluted 50% in culture medium supplemented with 8μgml−1 polybrene (Sigma) and immediately applied onto MutuDC1 cells. The plate was centrifuged for 60min at 2,000rpm at room temperature. 16h after spinfection the medium was exchanged for fresh complete IMDM-Glutamax medium. On day 4 after spinfection the lentiviral transduced MutuDC1 were isolated using FACs sorting.

### Mice

Xcr1^Cre-mTFP1B6^ mice were generated in-house by Malissen and colleagues and have been described previously ^4^. The 129S2-Ifnar1^tm1Agt^/Mmjax and the C57Bl/6-Tg(TcraTcrb)1100Mb/J (OT-I) mice (TheJacksonLaboratory) were provided by V. Andreakos (BRFAA). The B6.129P2-Gt(ROSA)26Sor^tm1(DTA)Lky^/J (Rosa26^LSL-DTA^) and the B6.SJL-Ptprc^a^ Pepc^b^/BoyJ (CD45.1) mice were originally purchased from TheJacksonLaboratory. All animal procedures were approved by the Veterinary Administration Bureau, Prefecture of Athens, Greece under compliance to the national law and the EU Directives and performed in accordance with the guidance of the Institutional Animal Care and Use Committee of BSRC Alexander Fleming.

### Syngeneic tumor models

Gender and age-matched, 6-12-week-old mice were used for all studies. Mice were housed with water and food ad libitum and 12-hour/12-hour night/daylight cycle under standard special pathogen-free conditions at BSRC Alexander Fleming and at the Centre d’immunologie Marseille Luminy. For all cancer models, cancer cells were thawed from frozen stocks and propagated in medium (RPMI + 10% FBS + 0.1% 2ME) for 5–7 days with one intervening passage in vitro. On the day of injection, cells were harvested by incubation in 0.05% trypsin-EDTA and washed three times with endotoxin-free PBS. For the orthotopic lung cancer models, mice were anesthetized via i.p. injection of xylazine and ketamine. LLC or CULA cells (2 × 10^5^) resuspended into 50 ul DMEM and enriched with 20% extracellular matrix (Matrigel, BD Biosciences), were intrapleurally injected to the left lung lobe of mice using a 29G needle (BD Biosciences). For the metastatic model, mice were injected intravenously via the tail vein with 5 x 10^5^ LLC cells in 100 μl DMEM. For the heterotopic cancer model 5 × 10^5^ LLC cells were resuspended in 100 μl of endotoxin-free PBS and injected subcutaneously on the left flank of C57BL/6 mice.

### Bone marrow chimeras

6–8-week-old C57BL/6 CD45.2 mice were lethally irradiated with 6.6Gy twice (3h rest in between exposures), then rested for 1d to recover. Bone marrow cells were prepared from the tibia and fibia bones of donor mice. Mice were injected i.v. with 2×10^6^ mixed BM cells (C57BL/6 CD45.1 : 129S2-Ifnar1^tm1Agt^/Mmjax CD45.2, 1:1 ratio) in a total volume of 0.2ml. After 8 weeks, chimerism was quantitated in the blood and mice were injected intravenously via the tail vein with 5 × 10^5^ LLC cells in 100 μl DMEM. After 16 days mice were euthanatized and tumors and spleens excised for further analysis.

### In vivo MutuDC1 transfer

MutuDC1 were treated with 20μM leupeptin hemisulfate (TOCRIS Cat. 1167) or PBS for 4h, washed and stained with Tag-it Violet™ Proliferation and Cell Tracking Dye (Biolegend Cat.425101) or CellTrace™ Far Red Cell Proliferation Kit (Invitrogen Cat.C34564). 1.5 x 10^6^ tagged leupeptin-treated MutuDC1 and 1.5 x 10^6^ cells tagged PBS-treated MutuDC1 were mixed, re-suspended in endotoxin-free PBS and injected intratumorally in flank-tumour bearing micer. 16 h after intratumoral injection tumors were excised isolated and analyzed by FACs. For therapeutic experiments, MutuDC1 were stimulated with 12.5 μg/ml Poly (I:C) (TOCRIS Cat. 4287) and 7.5 μg/ml R848 (Invivogen Cat. tlrl-848) for 16h, plus 20μM Leupeptin hemisulfate or endotoxin-free PBS for an additional 4h. PBS and leupeptin-treated MutuDC1 were collected and resuspended in array in endotoxin-free PBS at 10^7^ cells/mL. 2×10^6^ cells were injected intratumorally in flank tumor bearing mice. Tumor growth was measured on day 3 and 5 post injection using a digital caliper.

For GBP2 co-adoptively transfer experiment MutuDC1-dCAS9^KRAB^ sgNT versus MutuDC1-dCAS9^KRAB^ sgGBP2 cells were stained with Tag-it Violet™ Proliferation and Cell Tracking Dye (Biolegend Cat.425101) or CellTrace™ Far Red Cell Proliferation Kit (Invitrogen Cat.C34564). 1.5 x 10^6^ cells tagged MutuDC1-dCAS9^KRAB^ sgNT and 1.5 x 106 cells tagged MutuDC1-dCAS9^KRAB^ sgGBP2 cells were mixed, re-suspended in endotoxin-free PBS and injected intratumorally in flank tumor bearing mice. 16 h after intratumoral injection tumors were excised and analyzed by FACs. For therapeutic experiments, MutuDC1-dCAS9^KRAB^ sgNT and MutuDC1-dCAS9^KRAB^ sgGBP2 were stimulated with 12.5 μg/ml Poly (I:C) and 7.5 μg/ml R848 for 16h, collected and re-suspended in array in endotoxin-free PBS at 10^7^ cells/mL. 2×10^6^ cells were injected intratumorally in flank tumor bearing mice. On the same day mice were treated via i.p injection with 2×10^6^ OTI CD8+ T cells and 20 mg/kg anti-PDL1 antibody (GoInVivo™ Purified anti-mouse CD274 (B7-H1, PD-L1) Antibody, Biolegend Cat. 124329). Tumor growth was measured on day 2 and 4 post-injection using a digital caliper.

For therapeutic experiments tumor volume stated was calculated as L × l 2, considering the longest diameter (L) and its perpendicular (l) for each tumor. On day 4 post intratumoral injection tumors and inguinal lymph nodes isolated and analyzed by FACs.

### Generation of Tissue Culture Media and MutuDC1 treatment

Lung tumor tissue and healthy lungs were chopped and fragments were cultured for 24h in RPMI supplemented with 10% heat-inactivated human serum (SigmaAldrich; H4522), 1% L-glutamine, 1% penicillin/streptomycin, 20 mM Hepes. 1mg of tissue fragments was cultured in 140 μl cultured medium. Tumor culture medium (TCM) and Healthy Culture Medium (HCM) were purified by centrifugation at 1000g for 10min at 4°C and diluted at 40% before being added to the MutuDC1 line. Tissue culture medium-treated MutuDC1 was analyzed at 24h.

### MutuDC1 cytokine, bafilomycin and leupeptin treatments

MutuDC1 cells were treated with mouse recombinant 20ng/ml IFN-β1 (Biolegend Cat.581302), mouse 80ng/ml IFN-α (Miltenyi Cat. 130-093-131), and mouse recombinant 10ng/ml IFN-γ (PEPROTech Cat.315-05). For mTORC1 activation cells were treated with 5nM Bafilomycin A1 (Invivogen Cat. tlrl-baf1). For inhibition of lysosomal proteases MutuDC1 cells pre-treated with 20μM leupeptin for 4h. After incubation cells were washed twice with PBS and then treated with 20 ng/ml IFN-β1. MutuDC1 was analyzed 16h post-treatments.

### Tissue processing for flow cytometry

Lungs were cleared from blood via intracardiac infusion of PBS. Tumor and lung tissue were cut into fragments and enzymatically digested in 10% FBS/HBSS (Gibco) using Collagenase IV (Sigma-Aldrich; 1 mg/ ml; cat. no. C7657), Dispase II (Roche; 1 mg/ml; cat. no. SCM133) and Dnase I (Sigma-Aldrich; 0.09 mg/ml; cat. no. DN25) for 45 min at 37°C with agitation. Digested tissues were passed through a 70-µm cell strainer.

### Flow cytometry

For FACS sorting experiments, debris was removed using Debris Removal Solution (Miltenyi xat. No. 130-109-398). Cells were washed with FACS buffer (2% FBS/PBS/1.5 mM EDTA), centrifuged, and resuspended in FACS buffer. Nonspecific binding was blocked by incubating cells anti-mouse CD16/32 Fc block (Biolegend; cat. no. 101310). Sytox-Green viability dye (Thermo Fisher Scientific; cat. no. S7020), Zombie NIR (Biolegend; cat. no. 423105), Zombie Violet (Biolegend; cat. no. 423113), DAPI solution (Thermo Fisher Scientific; cat. no. 62248) or BD Horizon Fixable Viabilility Stain 700 (BD Pharmigen; cat. no. 564997) were used to exclude dead cells. Murine cells were stained for 30 min at 4°C with the following antibodies (all from Biolegend, unless otherwise stated): CD3 APC(Clone:17A2 Cat:100236) or PE(Clone:17A2 Cat:100205) or FITC(Clone:17A2 Cat:100203), B220 APC(Clone:RA3-6B2 Cat:103211) or PE(Clone:RA3-6B2 Cat:103207), NK1.1 APC(Clone:7H1 Cat:143011) or PE(Clone:7H1 Cat:143003), Ly6G APC(Clone:1A8 Cat:127613) or PE(Clone:1A8 Cat:127607) or PE(Clone:RA3-6B2 Cat:103207), NK1.1 APC(Clone:7H1 Cat:143011) or PE(Clone:7H1 Cat:143003), Ly6G APC(Clone:1A8 Cat:127613) or PE(Clone:1A8 Cat:127607) or BV605(Clone:1A8 Cat:127639), Ly6C APC(Clone:HK1.4 Cat:128015) or PE(Clone:HK1.4 Cat:128007), CD11b APC(Clone:M1/70 Cat:101211) or PE(Clone:M1/70 Cat:101207) or FITC(Clone:M1/70 Cat:101205) or BV510(Clone:M1/70 Cat:101245), CD49b PE(Clone:DX5 Cat:108907), SinglecF APC(Clone:S17007L Cat:155507), CD45 ALEXA 700(Clone:13/2.3 Cat:147716) or APC-Cy7(Clone:30-F11 Cat:103115) or BV 785(Clone:30-F11 Cat:103149), CD11c PECy7(Clone:N418 Cat:117317) or FITC(Clone:N418 Cat:117305), MHCII APC-Cy7(Clone:M5/114.15.2 Cat:107627) or PE(Clone:M5/114.15.2 Cat:107607), XCR1 BV650(Clone:ZET Cat:148220) or PE(Clone:ZET Cat:148203), F4/80 PE(Clone:BM8 Cat:123109), IFNAR1 PE(Clone:MAR1-5A3 Cat:127311), CD4 FITC(Clone:GK1.5 Cat:100405) or PECy7(Clone:GK1.5 Cat:100421), CD8 APC-Cy7(Clone:53-6.7 Cat:100713) or ALEXA 700(Clone:53-6.7 Cat:100729), CD25 PE(Clone:PC61.5 Cat:102007), FOXP3 APC(Clone:MF-14 Cat:126407), CD45.1 ALEXA 700(Clone:A20 Cat:110723) or FITC(Clone:A20 Cat:110705), CD45.2 BV785(Clone:104 Cat:109839) or FITC(Clone:104 Cat:109805). Tetramers SIINFEKL-PE and SIINFEKL-APC were provided by the National Institute of Health. Primary murine cDCs1 from lung tumors and healthy lungs were analyzed or sorted as XCR1^high^ CD45+Lin-(Lin:CD3, B220, NK1.1, CD11b, LY6C,LY6G, Ter119) CD11c^+^MHCII^+^. For Annexin V/propidium iodide (PI) stain, up to 2 × 10^6^ cells were resuspended in Binding Buffer (0.01 M Hepes, pH 7.4, 0.14 M NaCl, and 2.5 mM CaCl2) and stained with Annexin V FITC (lot# B284572, cat. no. 640906) and antibodies for 30 min at 4°C. Cells were washed and resuspended in PI– Binding Buffer solution (0.1 mg/ml; Sigma-Aldrich; cat. no. P4170). For intracellular staining, cells were fixed and permeabilized using the Intracellular Fixation & Permeabilization Buffer Set (eBioscience; cat. no. 88–8823-88), followed by staining with Gpnmb eFluor 660 and CCR7 APC for 30 min at 4°C. For intracellular staining of phosphorylated proteins, cells were fixed and permeabilized using the Intracellular Fixation & Permeabilization Buffer Set (eBioscience; cat. no. 88–8823-88) according to manufacturer’s recommendations for detection of intracellular phosphorylated proteins, followed by staining with pMTOR-PE-Cy7 (eBiosciences; cat. no. 25–9718-42) and pAKT for 1h at 4°C. Data were acquired on BD FACSCANTO II or BD FACSCelesta (BD Biosciences) and analysed using FlowJo (Tree Star, v9.9.6) or cells were sorted on FACSARIA III (BD Biosciences). Data were presented using Prism (Graphpad, v9).

### Ex-vivo assays

*Acidification assay.* 5×10^6^ digested tissue cells were resuspended in 1ml DMEM, containing 10% heat-inactivated FBS, 1% L-glutamine, 1% penicillin/streptomycin (Gibco) and 50nM Lysotracker Green DND-26 (Invitrogen cat. no. L7526) for 1h at 37°C. Cells were washed, stained with selected antibodies and analysed by FACS.

*Proteolysis assay.* 10^7^ digested tissue cells were cultured in 3ml DMEM, containing 10% heat-inactivated FBS, 1% L-glutamine, 1% penicillin/streptomycin (Gibco) and 25ug/ml DQ™ Green BSA (Invitrogen cat. no. D12050) for 16h. Cells were washed, stained with selected antibodies and analyzed by FACs.

*Apoptotic cancer cell phagocytosis assay*. Apoptotic LLC cells were prepared by three rapid freeze/thaw cycles (–140 oC /37oC). Primary murine cDCs1 were FACS-sorted from lung tumors and healthy lungs, cultured in 384 well plates in DC1 culture medium [DMEM 10% FBS, 1% L-glutamine, 1% sodium pyruvate, 1% MEM-NEAA, 1% penicillin/streptomycin, 55 uM 2-mercaptoethanol and 20 ng/ml of recombinant murine granulocyte-macrophage colony-stimulating factor (PeproTech, Rocky Hill, NJ, United States)] at 1:5 apoptotic cancer cell to cDC1 ratio, washed and analyzed by FACs at 24h, 48h, 72h, 96h.

*Endocytosis assay.* Primary murine cDCs1 were FACS-sorted from digested tissues, cultured in 384 well plates with DC1 culture medium in the presence of 50 μg/ml Dextran 488 (40 kDa) (ThermoFisher, Cat No. FD40S), washed and analyzed by FACs at 2h, 24h and 48h.

### Adoptive transfer of OTI T

Spleens were isolated from OT-I-tg mice and were forced to pass through 100-µm and 70-µm cell strainers. Blood cells were removed using eBioscience™ 1XRBC Lysis Buffer. Untouched OTI CD8^+^ T cells were negatively selected from peripheral blood mononuclear cells using anti-phycoerythrin (PE) MicroBeads (Miltenyi Biotech) after incubation with the following PE-conjugated antibodies: CD11b, CD11c, MHCII, B220, Ly6G, Ly6C, NK1.1 and CD4. For adoptive transfer experiments, OTI CD8^+^ T cells were resuspended in endotoxin-free PBS at 10^7^ cells/mL and 2×10^6^ cells were injected intraperitoneally.

### Human blood cDC1 purification and culture

Peripheral blood mononuclear cells (PMBCs) were obtained by gradient centrifugation using Ficoll-Paque (StemCell; cat. no. 07861). Untouched cDCs1 cells were stained with CD3 PE(Clone:UCHT1 Cat:980008), CD14 PE(Clone:63D3 Cat:367103), CD15 PE(Clone:HI98 Cat:301905), CD16 PE(Clone:3G8 Cat:302007), CD19 PE(Clone:HIB/9 Cat:302207), CD34 PE(Clone:561 Cat:343605), CD36 PE(Clone:5-271 Cat:336205), CD56 PE(Clone:5.1H11 Cat:362507), CD123 PE(Clone:6H6 Cat:306005), CD253 PE(Clone:RIK-2 Cat:308205), CD1c PE(Clone:L161 Cat:331505), CD45 ALEXA700(Clone:H130 Cat:304023), CD11c APC-CY7(Clone:Bu15 Cat:337217), panHLA APC(Clone:Tu39 Cat:361713), CD141 BV605(Clone:M80 Cat:344117) in the presence of human anti-Fc Receptor antibodies (TrueStain; Biolegend; cat. no. 101320). cDCs1 were FACS-sorted as CD45^+^Lin^-^ (Lin:CD3, CD14, CD15, CD16, CD19, CD34, CD36, CD56, CD123, CD253, CD1c) CD11c^+^, panHLA^+^, CD141^+^ and cultured in DMEM supplemented with 10% heat-inactivated human serum (SigmaAldrich; H4522), 1% L-glutamine, 1% penicillin/streptomycin, 1% sodium pyruvate, 1% MEM nonessential amino acids, 2 mM Hepes, β-mercaptoethanol and 20ng/ml huIFNβ in 384 well plates. After 16h cells were stained with 50nM Lysotracker Green DND-26 (Invitrogen cat. no. L7526) for 1h and analyzed by FACs

### Immunofluorescence stainings

#### Immunofluorescence stainings in primary cDCs1

5.000-10.000 FACS-sorted primary tissue cDC1 cells were cytospun at 400 g for 4 min. Slides were let dry for 5–10 min and fixed for 15 min using 4% paraformaldehyde. After washing twice with PBS 1× and permeabilizing for 10 min using 0.1% Triton-X/PBS, blocking was performed using 1% BSA in 0.1% Triton-X 100/PBS. Cells were stained overnight at 4°C using Lamp2 mouse monoclonal antibody (Novus Biologicals, Cat# NBP2-22217, RRID:AB_2722697)(1/200), GPNMB monoclonal antibody (CTSREVL), (eFluor 660, eBioscience™, Cat. 50-5708-82) (1/50), CD208/DC-LAMP rat monoclonal antibody (eurobio Cat.DDX0192P-100) (1/100), Galectin 3 rabbit antibody (abcam, Cat. ab76245) (1/100). After washing, sections were incubated with secondary antibody Goat anti-Mouse IgG (H+L) Cross-Adsorbed Secondary Antibody, Alexa Fluor 555 (Thermo Fisher Scientific, catalog # A-21422, RRID AB_2535844) (1/500), Goat anti-Rabbit IgG (H+L) Cross-Adsorbed Secondary Antibody, Alexa Fluor 647 (Thermo Fisher Scientific, Cat. A21244)(1/500), Goat anti-Rat IgG (H+L) Cross-Adsorbed Secondary Antibody, Alexa Fluor 647 (Thermo Fisher Scientific, Cat. A21247) (1/500). Coverslips were mounted onto glass slides using DAPI Mounting Medium (Ibidi Cat.50011). Images were acquired using a TCS SP8X confocal system (Leica). Image analysis was performed using Leica LASX or Fiji-ImageJ.

#### Immunofluorescence stainings in MutuDC1

10^5^ MutuDC1 cells were seeded on Gelatin (Sigma Cat.SLCF9509) coated glass coverslips (VWR Cat.631-0170) in 24-well plates and fixed for 15 min using 4% paraformaldehyde. Cells were washed twice with PBS 1× and permeabilized for 10 min using 0.1% Triton-X/PBS. Blocking was performed using 1% BSA in 0.1% Triton-X 100/PBS. The primary antibody incubation was performed by inverting the coverslips onto 80–100-μl droplets of blocking buffer containing a dilution of the primary antibody for 1h at room temperature. The primary antibodies that have used was GBP 1-5, mouse Monoclonal Antibody (Santa Cruz, G-12): sc-166960) (1/300), Galectin 3 rabbit antibody (abcam, #Cat. ab76245) (1/100), CD208/DC-LAMP rat monoclonal antibody (eurobio #Cat.DDX0192P-100) (1/100) and Cathepsin B rabbit antibody (Cell Signaling Cat.D1C7Y)(1/300) The coverslips were then flipped back into the tissue culture plate and washed 3× with blocking buffer. The secondary antibodies were diluted in blocking buffer and added directly to the plate and incubated for 1h at room temperature. The secondary antibodies that have used was Goat anti-Mouse IgG (H+L) Cross-Adsorbed Secondary Antibody, Alexa Fluor 555 (from Thermo Fisher Scientific, catalog # A-21422, RRID AB_2535844)(1/500), Goat anti-Rabbit IgG (H+L) Cross-Adsorbed Secondary Antibody, Alexa Fluor 647 (from Thermo Fisher Scientific, catalog #Cat. A21244)(1/500), Goat anti-Rat IgG (H+L) Cross-Adsorbed Secondary Antibody, Alexa Fluor 647 (from Thermo Fisher Scientific, Cat.catalog # A21247)(1/500). Coverslips were mounted onto glass slides using DAPI Mounting Medium (Ibidi #Cat.50011). Images were acquired using a TCS SP8X confocal system (Leica). Image analysis was performed using Leica LASX or Fiji-ImageJ.

### Immuno-Electron microscopy

2×106 MutuDC1 cells were treated with 20 ng/ml murine recombinant IFN-β1 for 16h, washed twice with PBS, fixed with fixation buffer (3.5% paraformaldehyde and 0.05% glutaraldehyde in 0.1 M phosphate buffer, pH 7.2.), stored at 4°C and transferred to Creative Bioarray Inc. for immune-electron microscopy (Immuno-EM). For immuno-EM, cells were washed twice with PBS and dehydrated by a series of increasing concentrations of ethanol: 50%, 70% and 90% (15 min each). A mixture of LR-White and 90% ethanol (1:1) was added to the cells, followed by 2h incubation under agitation. After 2h the mixture was replaced by 100% embedding LR-White and incubated overnight at 4°C. The samples were transferred to gelatin capsules filled with LR-White and polymerized at 55oC for 24h. LR-white blocks were cut at 60-90 mm sections on a nickel grid (250 mesh). Sections were incubated with 0.05 M Glycine in PBS buffer for 20 min and blocked with 1% BSA in PBS for 30 min at room temperature. Sections were incubated with primary antibody overnight at 4°C using GBP 1-5, mouse Monoclonal Antibody (Santa Cruz, G-12): sc-166960). After 3 washes with PBS sections were incubated with 50 ul colloidal gold-IgG complex (10 nm) diluted in blocking buffer (1:20) for 30 min at 37°C. Sections were washed with PBS, incubated with aqueous uranyl acetate saturate solution for 1 min and analyzed by TEM.

### quantitative PCR

Total RNA was extracted from MutuDC1 using the Single Cell RNA Purification Kit (Norgen Biotek) according to the manufacturer’s instructions. Superscript II reverse-transcriptase (Thermo Fisher Scientific) was used for cDNA synthesis and SYBR Green (Thermo Fisher Scientific) for qPCR performed on the CFX96 Touch Real-Time PCR Detection System from BioRad. Transcript levels of GBP2 were determined relative to B2M reference gene, using the ΔΔCt method. The following primer sets were used: GBP2 forward (Fw): 5’-TTCCCGGGTTACTACAGGGT -3’ and reverse (Rv): 5’- CTGGTCAGCTTTGCCTCTGA -3’ and murine B2M Fw: 5’- TTCTGGTGCTTGTCTCACTGA-3’and Rv: 5’- CAGTATGTTCGGCTTCCCATTC-3’.

### Murine bulk RNAseq analysis

#### cDNA Library preparation and FASTQ generation

Based on final cellular yield, RNA isolation and cDNA amplification was performed using either Smart-Seq® v4 Ultra® Low Input RNA Kit (Takara Clontech, cat# 635015) or NORGEN BIOTEK (cat# 51800). Prior to tagmentation, cDNA samples were loaded on a DNA High Sensitivity Chip on the 2100 Bioanalyzer (Agilent) to ensure transcript integrity, purity, and amount. RNA-seq libraries were subsequently constructed and sequenced on the Illumina NextSeq 2000 sequencer using Nextera XT or on the Ion Torrent PROTON sequencer using NEB Next Ultra RNA Library Prep Kit. Quality of FASTQ files, was assessed using FastQC (https://www.bioinformatics. babraham.ac.uk/projects/fastqc/) following the software recommendations. Bam files containing reads that were uniquely aligned to the murine were summarized to read counts table using the featureCounts package ^5^. Samples that did not reach quality standards were considered outliers and were omitted from the analysis.

The resulting gene counts table was subjected to DEA with metaseqr2 function, using DESeq2 algorithm for the normalization and statistical testing [Love et al. 2014]. Genes with less than five counts in 75% of the samples were excluded from downstream analysis. DEA thresholds were set for FDR equal to 0.01 and for logFC +/− 1. DAVID analysis was performed as previously described ^6^. GO using DAVID was performed for terms classified as biological processes.

#### Network Analysis

Gene regulatory network (GRN) inference from upregulated genes with FDR less than 0.05 was performed using GENIE3^7^. TF enrichment analysis in a curated gene set (Supplementary Table 1) was performed using ChIP-X Enrichment Analysis (Chea3) ^8^. RcisTarget was used to identify over-represented TF binding motifs (TFBS) with Normalized Enrichment Score (NES) >3. As inputs for RcisTarget, the curated gene set and the provided “mm9-tss-centered-10kb” database with genome-wide cross species rankings for each motif were used. For motif annotation to TFs, the default TF annotation dataset “motifAnnotations_mgi” was applied. TFs annotated to motifs were divided into high-confidence TFs, produced by Orthology or direct annotation, and low-confidence TFs mainly produced by motif similarity.

### Murine single cell RNA-seq data analysis

Fastqs of scRNAseq data of wild type tumor and wild type healthy lung cDCs were downloaded from GEO accession GSE131957 with the accession numbers GSM3832735 and GSM3832737 and merged ^9^. Ambient mRNA was identified with *soupX* ^10^, followed by QC and filtering (max counts 15000, min counts 500, min cells 3, max mt percentage 20%). The resultant 4568 high quality single cell transcriptomes were normalized with computeSumFactors. Cell cycle and mitochondrial scores were regressed out and highly variable genes per batch calculated and used for PCA analysis. Louvain clustering and UMAP were performed based on the first 35 PCA components. Cell annotation was done using the markers produced with the Wilcoxon test. In the DC1 subcluster, we used the Seurat pipeline ^11^ i.e. normalization, high variable genes, scaling, PCA, neighborhood graph with 35 PCs, UMAP with umap-learn package, and Louvain clustering with resolution 5. Neighboring clusters with healthy-tumor ratio below 1 were merged to subpopulation “Tumor/C1”, and three other ∼mostly∼ healthy subpopulations occurred. Differentially expressed genes were calculated with the Seurat’s FindMarkers function with default parameters and the upregulated genes (FDR<0.05) were used as input for the DAVID functional annotation tool ^6^, using the *Mus musculus* genome as background. To infer GRNs pySCENIC analysis was performed ^12^. pySCENIC consisted of generation of co-expression modules with GRNBoost2, refinement of modules with RcisTarget and evaluation of regulon activity with AUCell ^12^. Regulons were incorporated as a new assay in a Seurat pipeline and the regulon specificity score was calculated for each cluster.

### Human single cell RNA-seq data analysis

scRNAseq meta-data of cDCs of human lung, liver, colon and stomach tumors and their paired healthy tissues were accessed from the FG-Lab Resources platform, maintained by gustaveroussy (https://gustaveroussy.github.io/FG-Lab/). We visualized the cluster density on UMAP plots with a customized script using ggplot2. Differentially expression analysis between tumor and healthy cDCs1 was performed using Wilcoxon rank sum test. Functional enrichment analysis was performed using the enrichR package ^13^. GRNswere inferred with pySCENIC as previously described ^12^.

### Statistics

Statistical analysis was performed using GraphPad Prism software version 8. No formal randomization was performed when comparisons were done across mice of different genotypes. Mice of the same genotypes receiving different treatments were randomized according to their tumor volume size. Investigators were blinded the genotype of the mice during sample collection and preparation. Investigators were not blinded during FACS experiments as all samples were analyzed equivalently within defined experimental groups.

### Reporting summary

Further information on research design is available in the Nature Research Reporting Summary linked to this paper.

### Data Availability

The RNA-seq data generated during the course of this study has been deposited and will be available on the GEO database. Publicly available human scRNAseq data can be accessed from the FG-Lab Resources platform, maintained by gustaveroussy (https://gustaveroussy.github.io/FG-Lab/). Murine scRNAseq data can be downloaded from GEO accession GSE131957 with the accession numbers GSM3832735 and GSM3832737. All the other primary data and materials are available from the corresponding author upon request.

## REFERENCES

1 Canton, J. et al. The receptor DNGR-1 signals for phagosomal rupture to promote cross-presentation of dead-cell-associated antigens. Nature immunology 22, 140–153 (2021). https://doi.org:10.1038/s41590-020-00824-x

2 Theisen, D. J. et al. WDFY4 is required for cross-presentation in response to viral and tumor antigens. Science 362, 694–699 (2018). https://doi.org:10.1126/science.aat5030

3 Ferris, S. T. et al. cDC1 prime and are licensed by CD4(+) T cells to induce anti-tumour immunity. Nature 584, 624–629 (2020). https://doi.org:10.1038/s41586-020-2611-3

4 Broz, M. L. et al. Dissecting the tumor myeloid compartment reveals rare activating antigen-presenting cells critical for T cell immunity. Cancer cell 26, 638–652 (2014). https://doi.org:10.1016/j.ccell.2014.09.007

5 Salmon, H. et al. Expansion and Activation of CD103(+) Dendritic Cell Progenitors at the Tumor Site Enhances Tumor Responses to Therapeutic PD-L1 and BRAF Inhibition. Immunity 44, 924–938 (2016). https://doi.org:10.1016/j.immuni.2016.03.012

6 Lavin, Y. et al. Innate Immune Landscape in Early Lung Adenocarcinoma by Paired Single-Cell Analyses. Cell 169, 750–765 e717 (2017). https://doi.org:10.1016/j.cell.2017.04.014

7 Bottcher, J. P. et al. NK Cells Stimulate Recruitment of cDC1 into the Tumor Microenvironment Promoting Cancer Immune Control. Cell 172, 1022–1037 e1014 (2018). https://doi.org:10.1016/j.cell.2018.01.004

8 Prokhnevska, N. et al. CD8(+) T cell activation in cancer comprises an initial activation phase in lymph nodes followed by effector differentiation within the tumor. Immunity 56, 107–124 e105 (2023). https://doi.org:10.1016/j.immuni.2022.12.002

9 Ngo, C. C. & Man, S. M. Mechanisms and functions of guanylate-binding proteins and related interferon-inducible GTPases: Roles in intracellular lysis of pathogens. Cellular microbiology 19 (2017). https://doi.org:10.1111/cmi.12791

10 Embgenbroich, M. & Burgdorf, S. Current Concepts of Antigen Cross-Presentation. Frontiers in immunology 9, 1643 (2018). https://doi.org:10.3389/fimmu.2018.01643

11 Zhang, S., Chopin, M. & Nutt, S. L. Type 1 conventional dendritic cells: ontogeny, function, and emerging roles in cancer immunotherapy. Trends in immunology 42, 1113–1127 (2021). https://doi.org:10.1016/j.it.2021.10.004

12 Bonam, S. R., Wang, F. & Muller, S. Lysosomes as a therapeutic target. Nature reviews. Drug discovery 18, 923–948 (2019). https://doi.org:10.1038/s41573-019-0036-1

13 Zhu, S. Y. et al. Lysosomal quality control of cell fate: a novel therapeutic target for human diseases. Cell death & disease 11, 817 (2020). https://doi.org:10.1038/s41419-020-03032-5

14 Murphy, T. L. & Murphy, K. M. Dendritic cells in cancer immunology. Cellular & molecular immunology 19, 3–13 (2022). https://doi.org:10.1038/s41423-021-00741-5

15 Wu, R. & Murphy, K. M. DCs at the center of help: Origins and evolution of the three-cell-type hypothesis. The Journal of experimental medicine 219 (2022). https://doi.org:10.1084/jem.20211519

16 Cabeza-Cabrerizo, M., Cardoso, A., Minutti, C. M., Pereira da Costa, M. & Reis e Sousa, C. Dendritic Cells Revisited. Annual review of immunology 39, 131–166 (2021). https://doi.org:10.1146/annurev-immunol-061020-053707

17 Kvedaraite, E. & Ginhoux, F. Human dendritic cells in cancer. Science immunology 7, eabm9409 (2022). https://doi.org:10.1126/sciimmunol.abm9409

18 Barry, K. C. et al. A natural killer-dendritic cell axis defines checkpoint therapy-responsive tumor microenvironments. Nature medicine 24, 1178–1191 (2018). https://doi.org:10.1038/s41591-018-0085-8

19 Michea, P. et al. Adjustment of dendritic cells to the breast-cancer microenvironment is subset specific. Nature immunology 19, 885–897 (2018). https://doi.org:10.1038/s41590-018-0145-8

20 Roberts, E. W. et al. Critical Role for CD103(+)/CD141(+) Dendritic Cells Bearing CCR7 for Tumor Antigen Trafficking and Priming of T Cell Immunity in Melanoma. Cancer cell 30, 324–336 (2016). https://doi.org:10.1016/j.ccell.2016.06.003

21 Ghislat, G. et al. NF-kappaB-dependent IRF1 activation programs cDC1 dendritic cells to drive antitumor immunity. Science immunology 6 (2021). https://doi.org:10.1126/sciimmunol.abg3570

22 Hildner, K. et al. Batf3 deficiency reveals a critical role for CD8alpha+ dendritic cells in cytotoxic T cell immunity. Science 322, 1097–1100 (2008). https://doi.org:10.1126/science.1164206

23 Garris, C. S. et al. Successful Anti-PD-1 Cancer Immunotherapy Requires T Cell-Dendritic Cell Crosstalk Involving the Cytokines IFN-gamma and IL-12. Immunity 49, 1148–1161 e1147 (2018). https://doi.org:10.1016/j.immuni.2018.09.024

24 Sanchez-Paulete, A. R. et al. Cancer Immunotherapy with Immunomodulatory Anti-CD137 and Anti-PD-1 Monoclonal Antibodies Requires BATF3-Dependent Dendritic Cells. Cancer discovery 6, 71–79 (2016). https://doi.org:10.1158/2159-8290.CD-15-0510

25 Lam, K. C. et al. Microbiota triggers STING-type I IFN-dependent monocyte reprogramming of the tumor microenvironment. Cell 184, 5338–5356 e5321 (2021). https://doi.org:10.1016/j.cell.2021.09.019

26 Hegde, S. et al. Dendritic Cell Paucity Leads to Dysfunctional Immune Surveillance in Pancreatic Cancer. Cancer cell 37, 289–307 e289 (2020). https://doi.org:10.1016/j.ccell.2020.02.008

27 Ruhland, M. K. et al. Visualizing Synaptic Transfer of Tumor Antigens among Dendritic Cells. Cancer cell 37, 786–799 e785 (2020). https://doi.org:10.1016/j.ccell.2020.05.002

28 Wohn, C. et al. Absence of MHC class II on cDC1 dendritic cells triggers fatal autoimmunity to a cross-presented self-antigen. Science immunology 5 (2020). https://doi.org:10.1126/sciimmunol.aba1896

29 Gutierrez-Martinez, E. et al. Cross-Presentation of Cell-Associated Antigens by MHC Class I in Dendritic Cell Subsets. Frontiers in immunology 6, 363 (2015). https://doi.org:10.3389/fimmu.2015.00363

30 Cohn, L. et al. Antigen delivery to early endosomes eliminates the superiority of human blood BDCA3+ dendritic cells at cross presentation. The Journal of experimental medicine 210, 1049–1063 (2013). https://doi.org:10.1084/jem.20121251

31 Wang, F., Gomez-Sintes, R. & Boya, P. Lysosomal membrane permeabilization and cell death. Traffic 19, 918–931 (2018). https://doi.org:10.1111/tra.12613

32 Koopmann, J. O. et al. Export of antigenic peptides from the endoplasmic reticulum intersects with retrograde protein translocation through the Sec61p channel. Immunity 13, 117–127 (2000). https://doi.org:10.1016/s1074-7613(00)00013-3

33 Meyer, M. A. et al. Breast and pancreatic cancer interrupt IRF8-dependent dendritic cell development to overcome immune surveillance. Nat Commun 9, 1250 (2018). https://doi.org:10.1038/s41467-018-03600-6

34 Spranger, S., Bao, R. & Gajewski, T. F. Melanoma-intrinsic beta-catenin signalling prevents anti-tumour immunity. Nature 523, 231–235 (2015). https://doi.org:10.1038/nature14404

35 Bottcher, J. P. & Reis e Sousa, C. The Role of Type 1 Conventional Dendritic Cells in Cancer Immunity. Trends in cancer 4, 784–792 (2018). https://doi.org:10.1016/j.trecan.2018.09.001

36 Schenkel, J. M. et al. Conventional type I dendritic cells maintain a reservoir of proliferative tumor-antigen specific TCF-1(+) CD8(+) T cells in tumor-draining lymph nodes. Immunity 54, 2338–2353 e2336 (2021). https://doi.org:10.1016/j.immuni.2021.08.026

37 Kerdidani, D. et al. Lung tumor MHCII immunity depends on in situ antigen presentation by fibroblasts. The Journal of experimental medicine 219 (2022). https://doi.org:10.1084/jem.20210815

38 Kerdidani, D. et al. Wnt1 silences chemokine genes in dendritic cells and induces adaptive immune resistance in lung adenocarcinoma. Nat Commun 10, 1405 (2019). https://doi.org:10.1038/s41467-019-09370-z

39 Cheng, S. et al. A pan-cancer single-cell transcriptional atlas of tumor infiltrating myeloid cells. Cell 184, 792–809 e723 (2021). https://doi.org:10.1016/j.cell.2021.01.010

40 Di Pilato, M. et al. CXCR6 positions cytotoxic T cells to receive critical survival signals in the tumor microenvironment. Cell 184, 4512–4530 e4522 (2021). https://doi.org:10.1016/j.cell.2021.07.015

41 Zilionis, R. et al. Single-Cell Transcriptomics of Human and Mouse Lung Cancers Reveals Conserved Myeloid Populations across Individuals and Species. Immunity 50, 1317–1334 e1310 (2019). https://doi.org:10.1016/j.immuni.2019.03.009

42 Maier, B. et al. A conserved dendritic-cell regulatory program limits antitumour immunity. Nature 580, 257–262 (2020). https://doi.org:10.1038/s41586-020-2134-y

43 Saade, M., Araujo de Souza, G., Scavone, C. & Kinoshita, P. F. The Role of GPNMB in Inflammation. Frontiers in immunology 12, 674739 (2021). https://doi.org:10.3389/fimmu.2021.674739

44 van der Lienden, M. J. C., Gaspar, P., Boot, R., Aerts, J. & van Eijk, M. Glycoprotein Non-Metastatic Protein B: An Emerging Biomarker for Lysosomal Dysfunction in Macrophages. International journal of molecular sciences 20 (2018). https://doi.org:10.3390/ijms20010066

45 Barral, D. C. et al. Current methods to analyze lysosome morphology, positioning, motility and function. Traffic 23, 238–269 (2022). https://doi.org:10.1111/tra.12839

46 Saftig, P. & Puertollano, R. How Lysosomes Sense, Integrate, and Cope with Stress. Trends in biochemical sciences 46, 97–112 (2021). https://doi.org:10.1016/j.tibs.2020.09.004

47 Saxton, R. A. & Sabatini, D. M. mTOR Signaling in Growth, Metabolism, and Disease. Cell 168, 960–976 (2017). https://doi.org:10.1016/j.cell.2017.02.004

48 Aibar, S. et al. SCENIC: single-cell regulatory network inference and clustering. Nature methods 14, 1083–1086 (2017). https://doi.org:10.1038/nmeth.4463

49 Diamond, M. S. et al. Type I interferon is selectively required by dendritic cells for immune rejection of tumors. The Journal of experimental medicine 208, 1989–2003 (2011). https://doi.org:10.1084/jem.20101158

50 Mattiuz, R. et al. Type 1 conventional dendritic cells and interferons are required for spontaneous CD4(+) and CD8(+) T-cell protective responses to breast cancer. Clinical & translational immunology 10, e1305 (2021). https://doi.org:10.1002/cti2.1305

51 Brombacher, E. C. & Everts, B. Shaping of Dendritic Cell Function by the Metabolic Micro-Environment. Frontiers in endocrinology 11, 555 (2020). https://doi.org:10.3389/fendo.2020.00555

52 Kaur, S. et al. Regulatory effects of mammalian target of rapamycin-activated pathways in type I and II interferon signaling. The Journal of biological chemistry 282, 1757–1768 (2007). https://doi.org:10.1074/jbc.M607365200

53 Su, X. et al. Interferon-gamma regulates cellular metabolism and mRNA translation to potentiate macrophage activation. Nature immunology 16, 838–849 (2015). https://doi.org:10.1038/ni.3205

54 Snyder, J. P. & Amiel, E. Regulation of Dendritic Cell Immune Function and Metabolism by Cellular Nutrient Sensor Mammalian Target of Rapamycin (mTOR). Frontiers in immunology 9, 3145 (2018). https://doi.org:10.3389/fimmu.2018.03145

55 Sukhbaatar, N., Hengstschlager, M. & Weichhart, T. mTOR-Mediated Regulation of Dendritic Cell Differentiation and Function. Trends in immunology 37, 778–789 (2016). https://doi.org:10.1016/j.it.2016.08.009

56 Huynh-Thu, V. A., Irrthum, A., Wehenkel, L. & Geurts, P. Inferring regulatory networks from expression data using tree-based methods. PloS one 5 (2010). https://doi.org:10.1371/journal.pone.0012776

57 Shlomovitz, I., Speir, M. & Gerlic, M. Flipping the dogma - phosphatidylserine in non-apoptotic cell death. Cell communication and signaling : CCS 17, 139 (2019). https://doi.org:10.1186/s12964-019-0437-0

58 Tan, J. X. & Finkel, T. A phosphoinositide signalling pathway mediates rapid lysosomal repair. Nature 609, 815–821 (2022). https://doi.org:10.1038/s41586-022-05164-4

59 Mulder, K. et al. Cross-tissue single-cell landscape of human monocytes and macrophages in health and disease. Immunity 54, 1883–1900 e1885 (2021). https://doi.org:10.1016/j.immuni.2021.07.007

60 Man, S. M., Place, D. E., Kuriakose, T. & Kanneganti, T. D. Interferon-inducible guanylate-binding proteins at the interface of cell-autonomous immunity and inflammasome activation. Journal of leukocyte biology 101, 143–150 (2017). https://doi.org:10.1189/jlb.4MR0516-223R

61 Papadopoulos, C., Kravic, B. & Meyer, H. Repair or Lysophagy: Dealing with Damaged Lysosomes. Journal of molecular biology 432, 231–239 (2020). https://doi.org:10.1016/j.jmb.2019.08.010

62 Skowyra, M. L., Schlesinger, P. H., Naismith, T. V. & Hanson, P. I. Triggered recruitment of ESCRT machinery promotes endolysosomal repair. Science 360 (2018). https://doi.org:10.1126/science.aar5078

63 Jia, J. et al. Galectin-3 Coordinates a Cellular System for Lysosomal Repair and Removal. Developmental cell 52, 69–87 e68 (2020). https://doi.org:10.1016/j.devcel.2019.10.025

64 Jia, J. et al. Galectins Control mTOR in Response to Endomembrane Damage. Molecular cell 70, 120–135 e128 (2018). https://doi.org:10.1016/j.molcel.2018.03.009

65 Zhang, H., Zoued, A., Liu, X., Sit, B. & Waldor, M. K. Type I interferon remodels lysosome function and modifies intestinal epithelial defense. Proceedings of the National Academy of Sciences of the United States of America 117, 29862–29871 (2020). https://doi.org:10.1073/pnas.2010723117

66 Bretou, M. et al. Lysosome signaling controls the migration of dendritic cells. Science immunology 2 (2017). https://doi.org:10.1126/sciimmunol.aak9573

67 Nakatani, T. et al. The lysosomal Ragulator complex plays an essential role in leukocyte trafficking by activating myosin II. Nat Commun 12, 3333 (2021). https://doi.org:10.1038/s41467-021-23654-3

68 Wandel, M. P. et al. GBPs Inhibit Motility of Shigella flexneri but Are Targeted for Degradation by the Bacterial Ubiquitin Ligase IpaH9.8. Cell host & microbe 22, 507–518 e505 (2017). https://doi.org:10.1016/j.chom.2017.09.007

69 Oliveira, M. M. S. & Westerberg, L. S. Cytoskeletal regulation of dendritic cells: An intricate balance between migration and presentation for tumor therapy. Journal of leukocyte biology 108, 1051–1065 (2020). https://doi.org:10.1002/JLB.1MR0520-014RR

70 Jacquelot, N. et al. Sustained Type I interferon signaling as a mechanism of resistance to PD-1 blockade. Cell research 29, 846–861 (2019). https://doi.org:10.1038/s41422-019-0224-x

71 Lukhele, S. et al. The transcription factor IRF2 drives interferon-mediated CD8(+) T cell exhaustion to restrict anti-tumor immunity. Immunity 55, 2369–2385 e2310 (2022). https://doi.org:10.1016/j.immuni.2022.10.020

72 Duong, E. et al. Type I interferon activates MHC class I-dressed CD11b(+) conventional dendritic cells to promote protective anti-tumor CD8(+) T cell immunity. Immunity 55,308–323 e309 (2022). https://doi.org:10.1016/j.immuni.2021.10.020

73 Schaupp, L. et al. Microbiota-Induced Type I Interferons Instruct a Poised Basal State of Dendritic Cells. Cell 181, 1080–1096 e1019 (2020). https://doi.org:10.1016/j.cell.2020.04.022

74 Ardouin, L. et al. Broad and Largely Concordant Molecular Changes Characterize Tolerogenic and Immunogenic Dendritic Cell Maturation in Thymus and Periphery. Immunity 45, 305–318 (2016). http://dx.doi.org/10.1016/j.immuni.2016.07.019

## METHOD REFERENCES

1 Kerdidani, D. et al. Wnt1 silences chemokine genes in dendritic cells and induces adaptive immune resistance in lung adenocarcinoma. Nature communications 10, 1405, doi:10.1038/s41467-019-09370-z (2019).

2 Kerdidani, D. et al. Lung tumor MHCII immunity depends on in situ antigen presentation by fibroblasts. The Journal of experimental medicine 219, doi:10.1084/jem.20210815 (2022).

3 Fuertes Marraco, S. A. et al. Novel murine dendritic cell lines: a powerful auxiliary tool for dendritic cell research. Frontiers in immunology 3, 331, doi:10.3389/fimmu.2012.00331 (2012).

4 Wohn, C. et al. Absence of MHC class II on cDC1 dendritic cells triggers fatal autoimmunity to a cross-presented self-antigen. Science immunology 5, doi:10.1126/sciimmunol.aba1896 (2020).

5 Liao, Y., Smyth, G. K. & Shi, W. featureCounts: an efficient general purpose program for assigning sequence reads to genomic features. Bioinformatics 30, 923–930, doi:10.1093/bioinformatics/btt656 (2014).

6 Huang, D. W. et al. The DAVID Gene Functional Classification Tool: a novel biological module-centric algorithm to functionally analyze large gene lists. Genome biology 8, R183, doi:10.1186/gb-2007-8-9-r183 (2007).

7 Huynh-Thu, V. A., Irrthum, A., Wehenkel, L. & Geurts, P. Inferring regulatory networks from expression data using tree-based methods. PloS one 5, doi:10.1371/journal.pone.0012776 (2010).

8 Keenan, A. B. et al. ChEA3: transcription factor enrichment analysis by orthogonal omics integration. Nucleic acids research 47, W212–W224, doi:10.1093/nar/gkz446 (2019).

9 Maier, B. et al. A conserved dendritic-cell regulatory program limits antitumour immunity. Nature 580, 257–262, doi:10.1038/s41586-020-2134-y (2020).

10 Young, M. D. & Behjati, S. SoupX removes ambient RNA contamination from droplet-based single-cell RNA sequencing data. GigaScience 9, doi:10.1093/gigascience/giaa151 (2020).

11 Hao, Y. et al. Integrated analysis of multimodal single-cell data. Cell 184, 3573–3587 e3529, doi:10.1016/j.cell.2021.04.048 (2021).

12 Van de Sande, B. et al. A scalable SCENIC workflow for single-cell gene regulatory network analysis. Nature protocols 15, 2247–2276, doi:10.1038/s41596-020-0336-2 (2020).

13 Chen, E. Y. et al. Enrichr: interactive and collaborative HTML5 gene list enrichment analysis tool. BMC bioinformatics 14, 128, doi:10.1186/1471-2105-14-128 (2013).

