## Supplementary Figures for "Interferon-induced lysosomal membrane permeabilization and death cause cDC1-deserts in tumors"

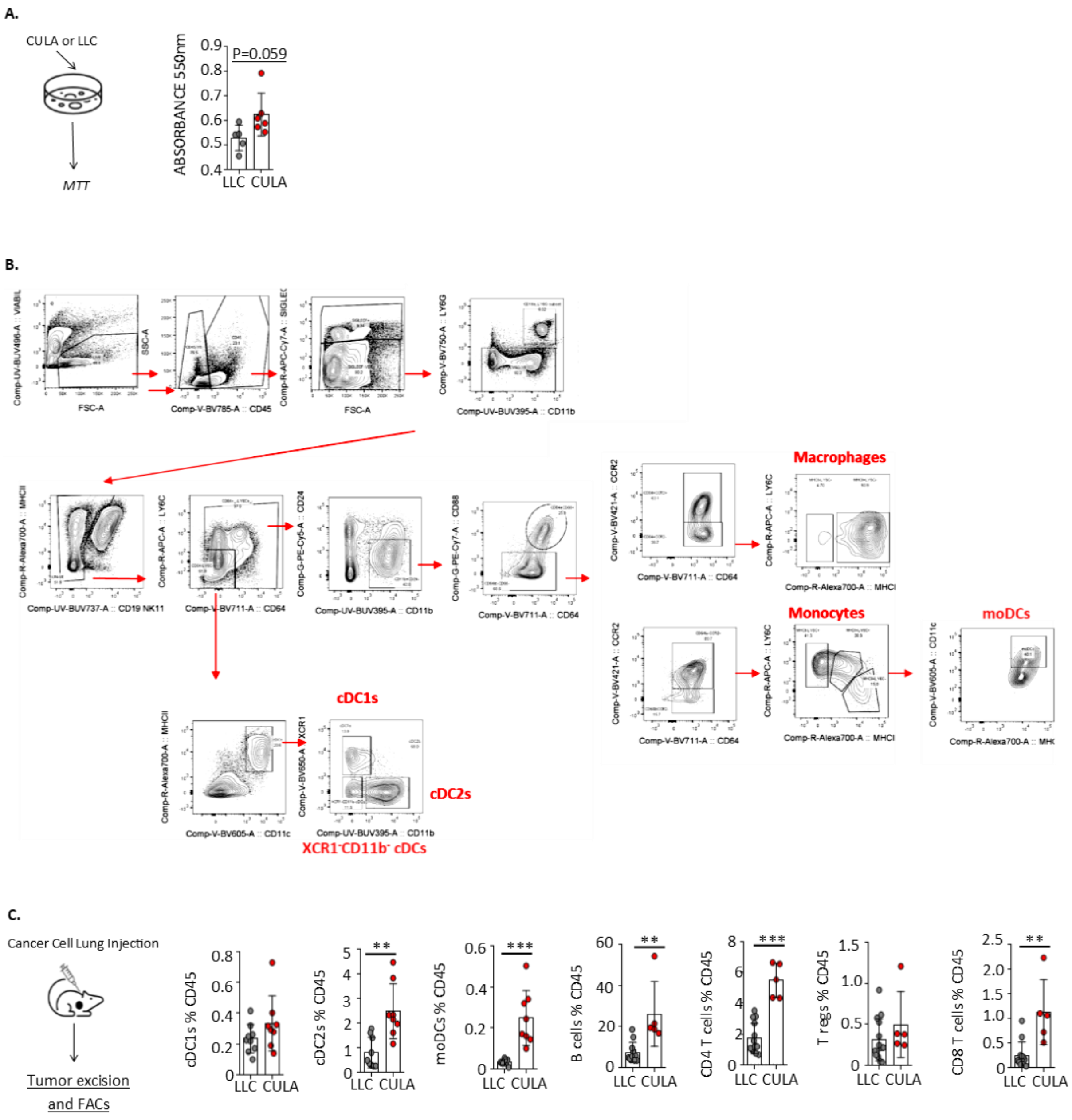


**Supplementary Figure 1. Increased immunogenicity of CULA relatively to LLC lung cancer cells in orthotopic lung cancer models. A.** In vitro proliferation rates based on the MTT assay. Cumulative data from 2 independent experiments with 3 technical replicates per cell line. **B.** Gating strategy to identify intratumoral cDC1, cDC2, monocyte-derived DCs, monocytes and macrophages. **C.** CULA versus LLC cancer cells were injected on the left lung lobe of immunocompetent syngeneic mice. Lung tumors were excised, digested and analyzed by FACS. Cumulative data from 2 independent experiments with 3-4 biological replicates per group per experiment. Error bars, mean± sem; *p<0.05, **p<0.01, ***<0.001, two-tailed unpaired t-test.

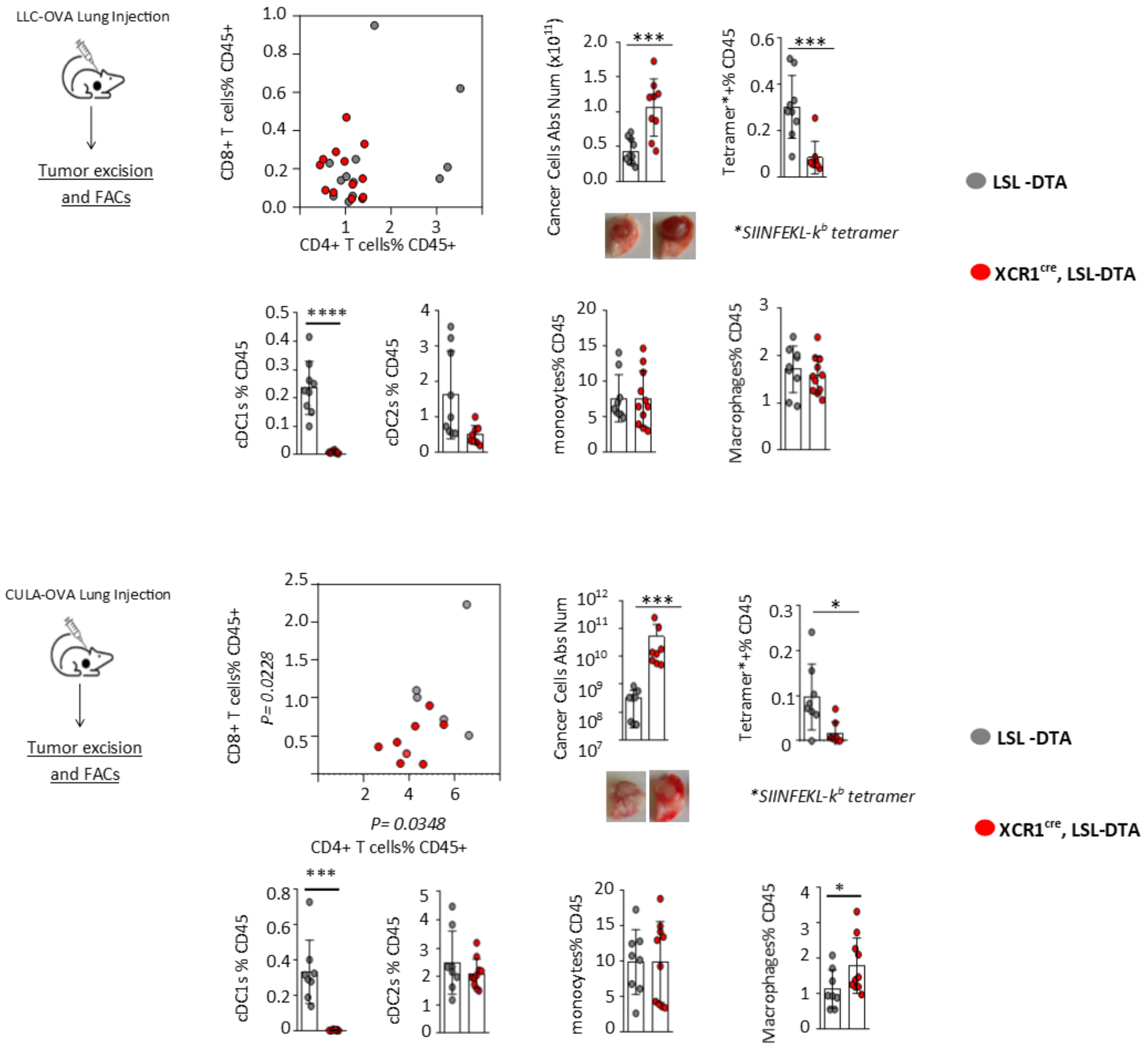


**Supplementary Figure 2. XCR1$^{cre}$ LSL-DTA mice display compromised lung cancer immunity.** The XCR1$^{cre}$ line was crossed to LSL-DTA to constitutively ablate cDC1s. LLC-OVA$^{mCherry}$ or the less aggressive more immunogenic CULA-OVA$^{mCherry}$ were injected in the left lung lobe of immunocompetent syngeneic mice. Lung tumors were excised on day 12 (LLC model) or day 28 (CULA model), digested, stained and analyzed by FACS. From left to right. Absolute number of mCherry cancer cells enumerated with FACS counting beads. OVA-specific CD8$^+$ T cells percent total CD45$^+$ T cells, identified with SIINFELK-k$^b$ tetramers. Myeloid cell subsets percent CD45$^+$ cells. Data are pooled from two independent experiments with 3-4 biological replicates per group per experiment. Error bars, mean± sem; *p<0.05, ***p<0.001, ****<0.0001, two-tailed unpaired t-test.

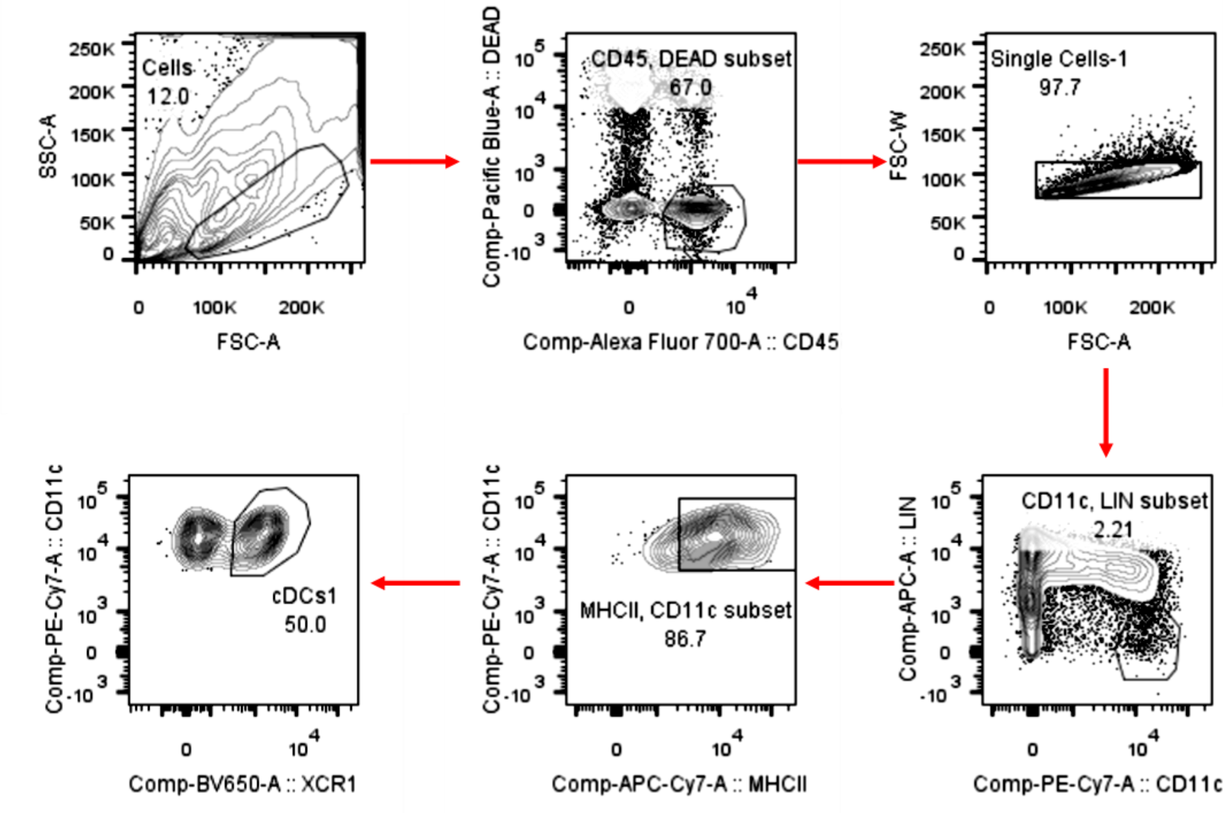


**Supplementary Figure 3. Identification of murine lung tumor cDC1 by FACS.** LLC cells was injected on the left lung lobe of mice. Lung tumors were excised, digested and analyzed by FACS. Gating strategy to identify cDC1 is shown.

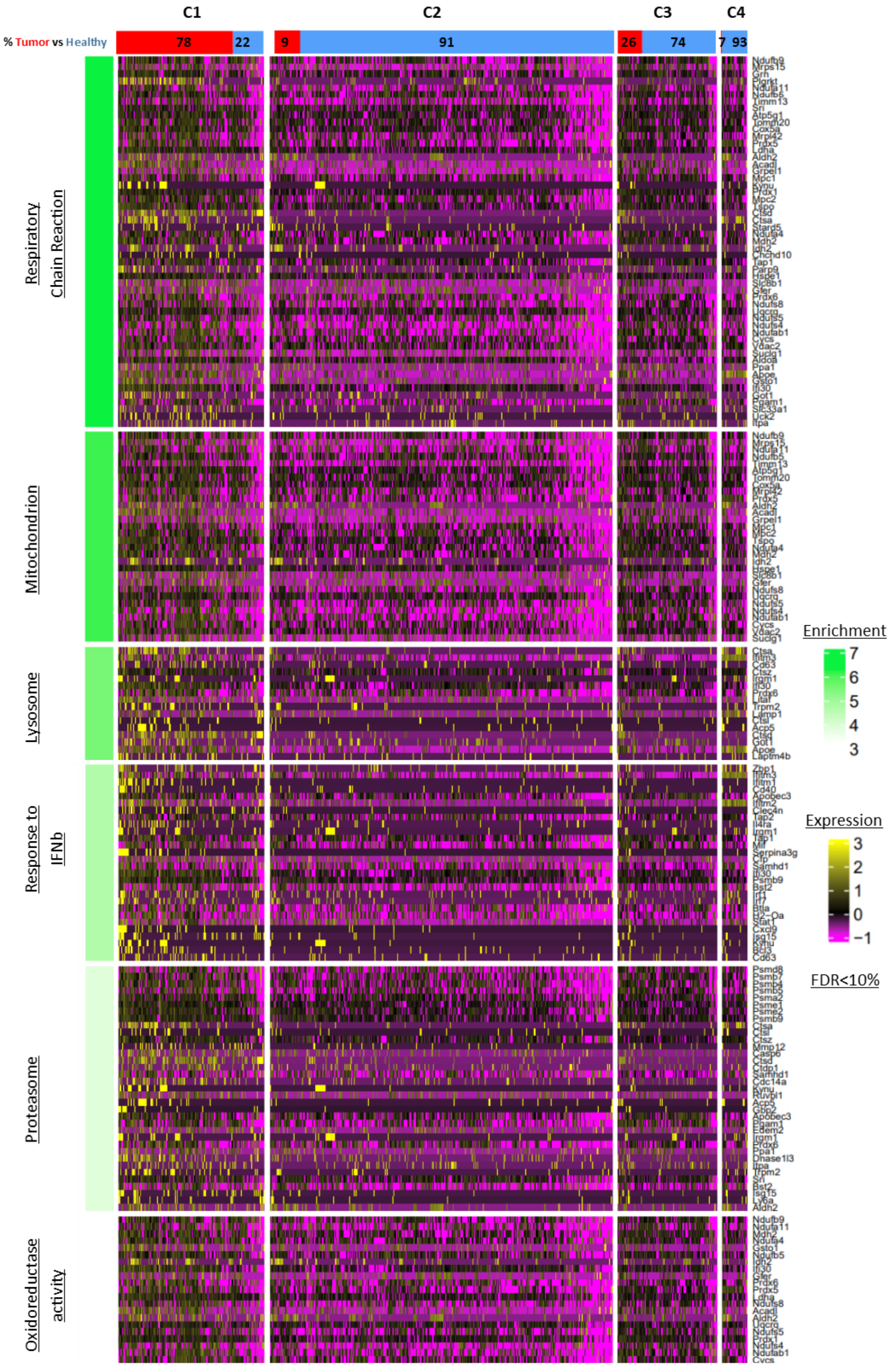
C1
C2
C3
C4
% Tumor vs Healthy
78
22
9
91
26
74
7
93
Respiratory Chain Reaction
Mitochondrion
Lysosome
Response to IFNb
Proteasome
Oxidoreductase activity
Enrichment
7
6
5
4
3
Expression
3
2
1
0
−1
FDR<10%

**Supplementary Figure 4. Lung tumour cDC1 transcriptomes are enriched in metabolic, lysosomal and type I IFN response pathways.** KP lung tumor and healthy lung cDC1 single cell transcriptomes were extracted from our (Merad and colleagues) public murine dataset [42]. Heatmap depicts deregulated genes that were found enriched in pathway enrichment analysis in murine cDC1 clusters.

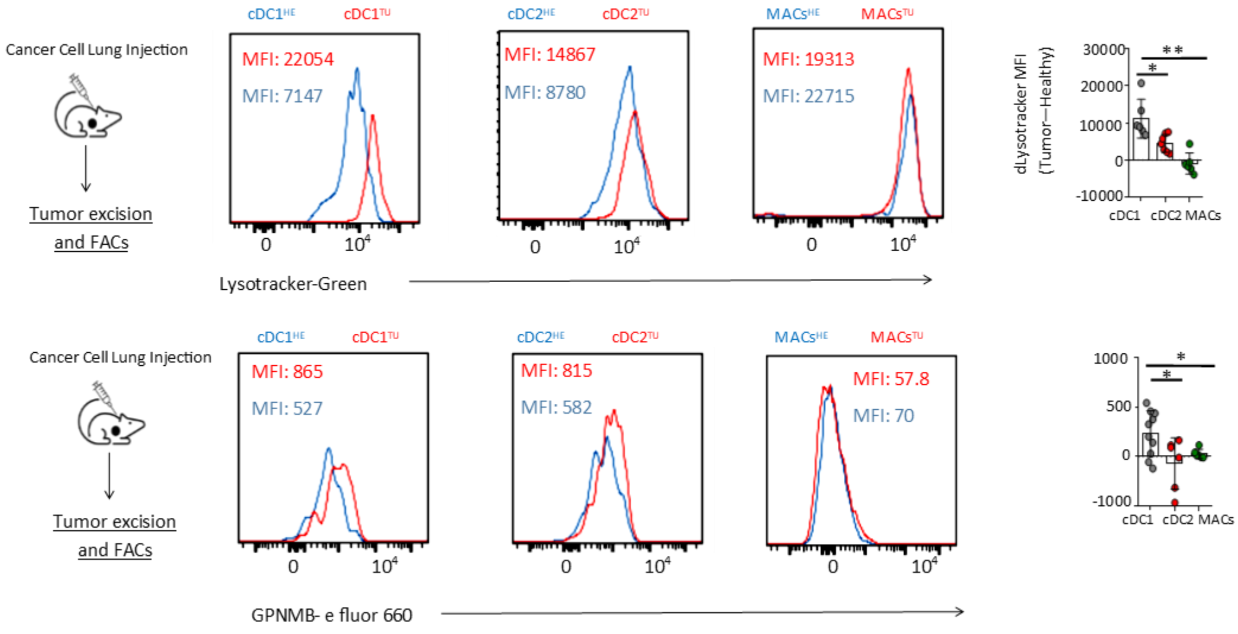


**Supplementary Figure 5. Increased lysosomal stress response in lung cancer cDCs1 relatively to cDCs2 and macrophages.** LLC-OVA$^{mCherry}$ were injected in the left lung lobe of immunocompetent syngeneic mice. Lung tumors were excised digested, stained and analyzed by FACS. **A.** Endo-lysosomal acidification assessed by Lysotracker stain. Representative histogram plots and cumulative data. **B.** Expression of the lysosome stress marker Gpnmb (intracellular). Representative histogram plots and cumulative data. **A-B.** Representative or cumulative data pooled from two independent experiments with 3-4 biological replicates per group per experiment. Error bars, mean± sem; *p<0.05, **p<0.01, two-tailed unpaired t-test.

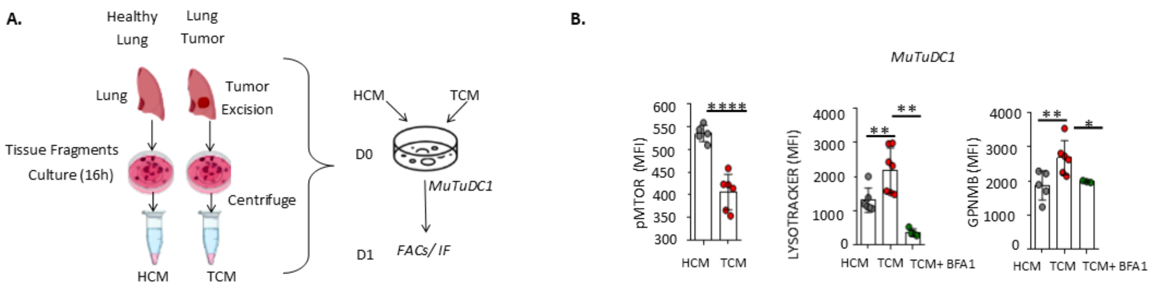


**Supplementary Figure 6. Tumour-induced lysosomal stress in MutuDC1 is suppressed by the mTOR activator Bafilomycin.** Tumor tissue and healthy lungs were chopped and fragments were cultured for 24h. Tumor Culture Medium (TCM) and Healthy tissue Culture Medium (HCM) were purified and added to the MutuDC1 line ($MutuDC1^{TCM}$ versus $MutuDC1^{HCM}$). After 16h FACS analysis was performed. From left to right. TCM-supressed phospho-mTOR and lysosomal stress (intracellular Gpnmb). Endo-lysosomal responses (Lysotracker) to mTOR activation (Bafilomycin) in the presence of TCM. Cumulative data from two independent experiments with 3-4 replicates per group per experiment. Error bars, mean± sem; *p<0.05, **p<0.01, ****p<0.0001, two-tailed unpaired t-test.

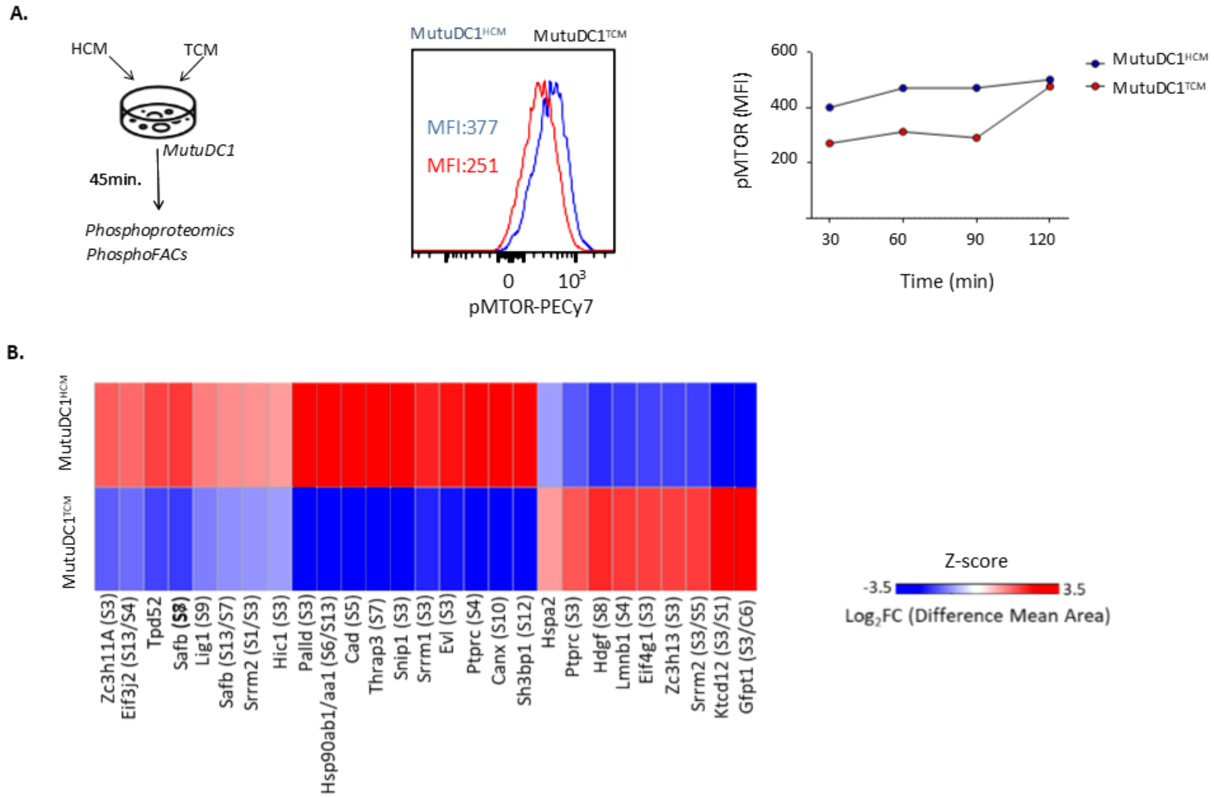


**Supplementary Figure 7. Phosphoproteomics analysis of Tumour Culture Medium-exposed MutuDC1$^{TCM}$ versus Healthy tissue Culture Medium-exposed MutuDC1$^{HCM}$.** **A.** Experimental scheme and kinetics of mTOR inhibition by TCM. **B.** Differential phosphorylation analysis-based heatmap at 90min. Cumulative data from one experiment with 2 replicates per group.

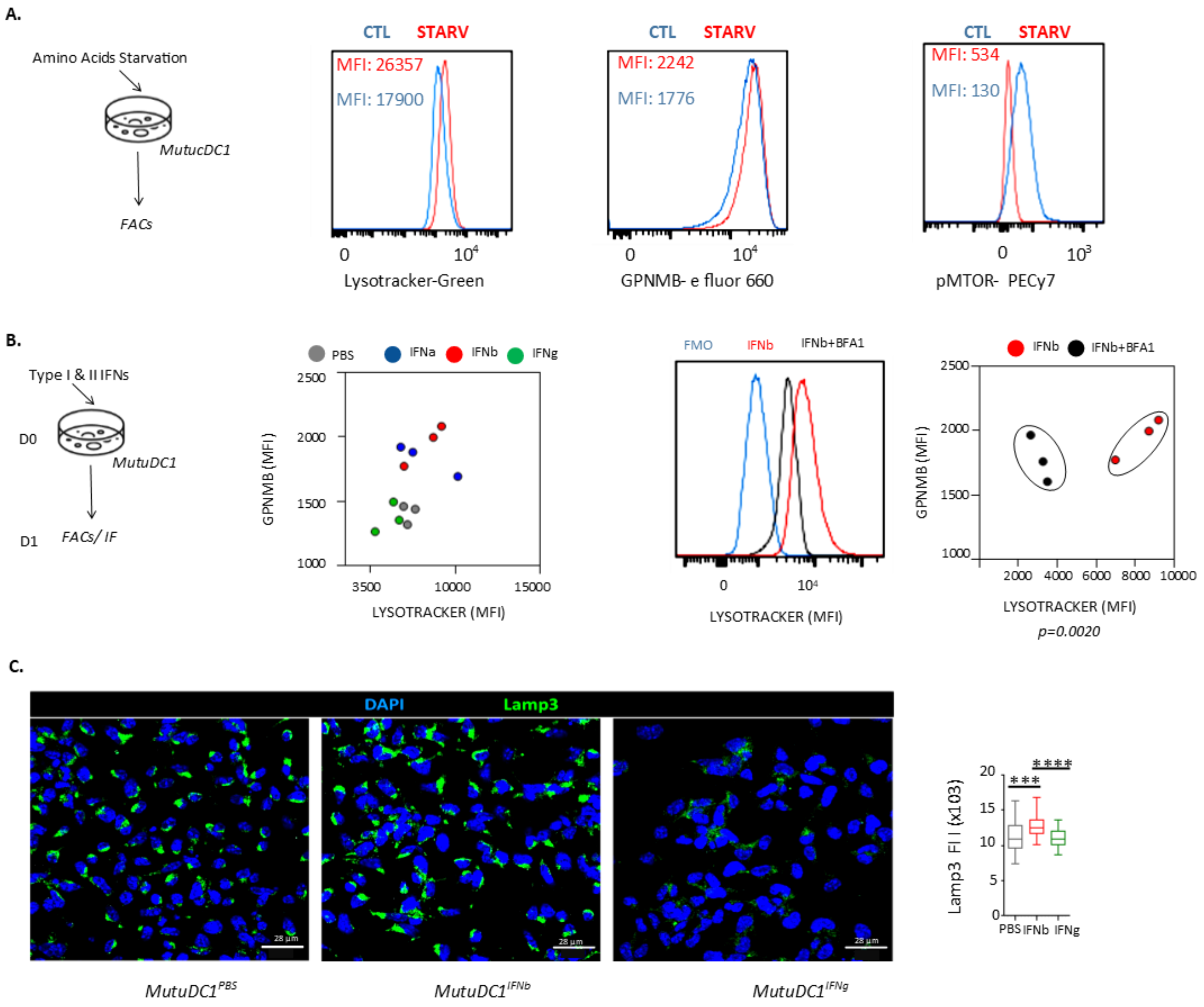


**Supplementary Figure 8. Amino acid starvation and type I IFNs induce lysosomal stress and lysosome biogenesis in MutuDC1. A.** MutuDC1 responses to amino acid deprivation. From left to right. Experimental scheme. Endo-lysosomal responses (Lysotracker), lysosomal stress (intracellular Gpnmb) and mTOR activation level. **B-C.** MutuDC1 responses to type I (IFNα, IFNβ) and type II IFN (IFNγ). **B.** From left to right. Experimental scheme. Endo-lysosomal responses (Lysotracker) and lysosomal stress (intracellular Gpnmb). Endo-lysosomal responses (Lysotracker) to mTOR activation (Bafilomycin) in the presence of IFNβ. Representative or cumulative data from two independent experiments with 3-4 replicates per group per experiment. **C.** Lamp3 quantification per cell by image analysis of the lysosomal membrane protein LAMP3 (n=40 cells per condition). Error bars, mean± sem; ***p<0.001, ****p<0.0001, two-tailed unpaired t-test.

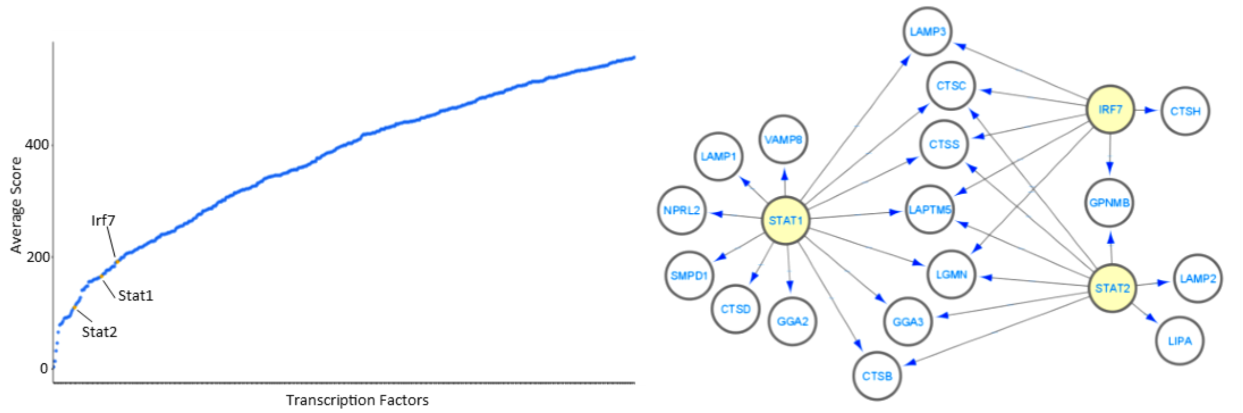


**Supplementary Figure 9. Transcription Factor Enrichment analysis predicts strong regulation of lysosomal genes by IFN-induced TFs.** Left graph depicts TF rank as produced by the Mean Rank of Chea3 of TFs that regulate a curated list of genes involved in lysosomal processes (Supplementary Tabe 1). Right. Representation of TF-gene actions via Cytoscape. Yellow nodes depict TFs, white nodes genes and arrows regulatory actions.

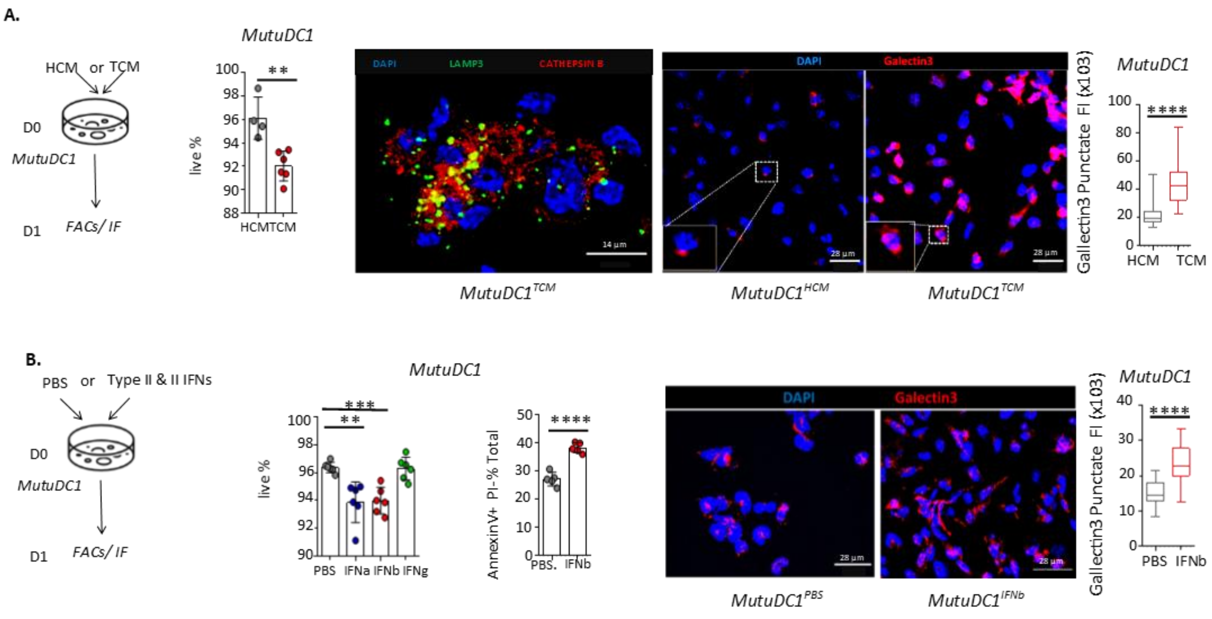

**Supplementary Figure 10. Tumour Culture Medium and Type I IFNs induces lysosomal membrane permeabilization and death in MutuDC1. A.** MutuDC1 responses to Tumour Culture Medium. From left to right. Experimental scheme. Percent live cells. Cathepsin B leakage from the lysosomes based on LAMP3/Cathepsin B co-stain. Galectin 3 puncta and quantification by image analysis (n=80 cells per condition). **B.** MutuDC1 responses to type I (IFNα, IFNβ) and type II IFN (IFNγ). From left to right. Experimental scheme. Percent live cells. Percent annexin V+ cells (pre-death state). Galectin 3 puncta and quantification by image analysis (n=21 cells per condition). Error bars, mean± sem; ***p<0.001, ****p<0.0001, two-tailed unpaired t-test.

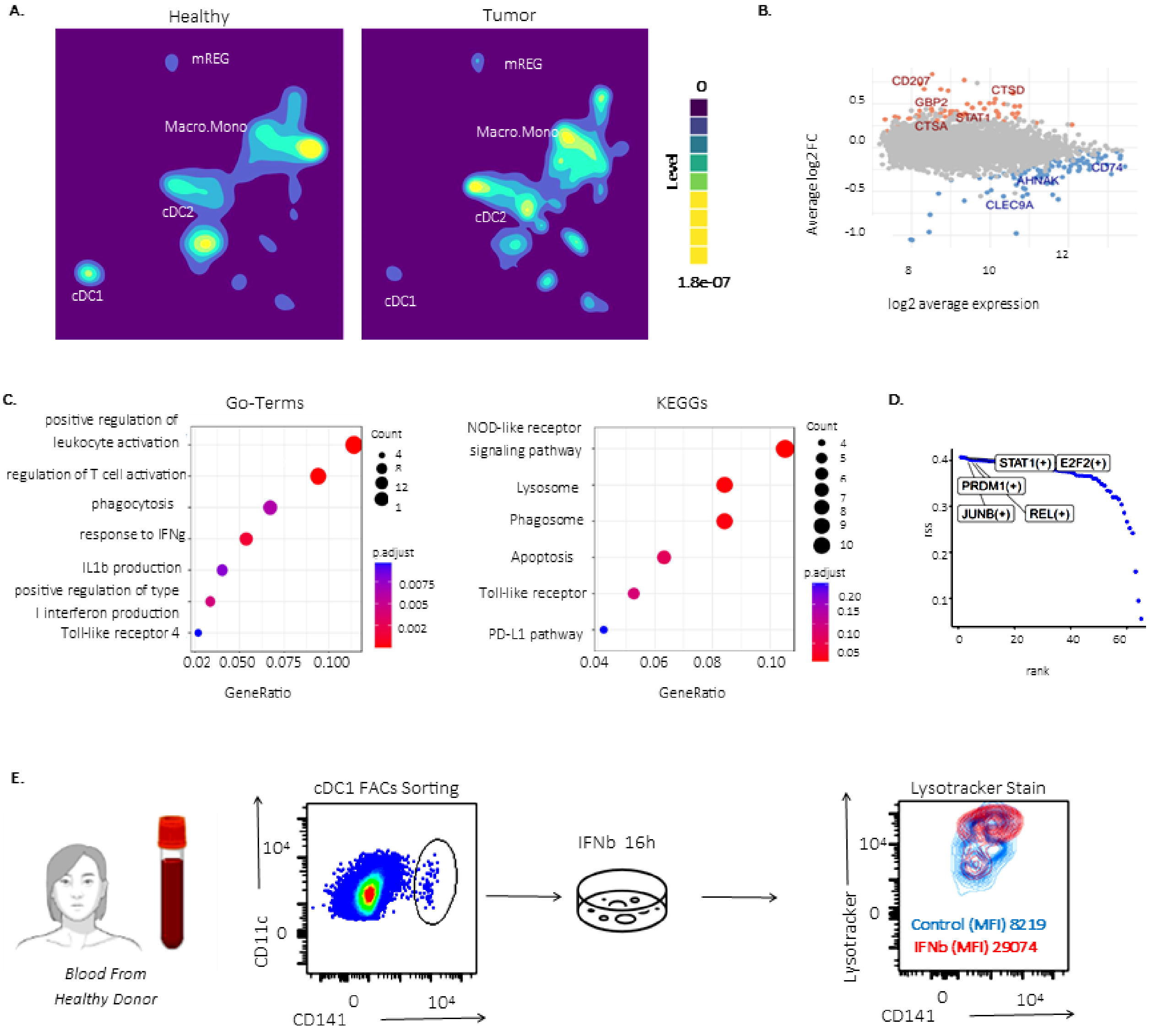

**Supplementary Figure 11. Evidence for IFN-driven lysosomal stress in human intratumoural cDCs1.** A publicly available scRNAseq dataset consisting of mononuclear phagocytes of paired tumor and healthy tissues (lung, liver, colon, stomach) was downloaded[59]. **A.** cDC subset density plots in tumour and healthy tissues. **B.** Mean-Average(MA)-plot of log2 average expression versus log2 fold change between human tumour cDC1 versus paired unaffected tissue cDC1 (public dataset). Colored dots indicate genes that are significantly differentially expressed. **C.** Dotplot of GO terms and KEGG pathways in pathway enrichment analysis. **D.** Ranking of tumor cDC1 regulons (SCENIC) according to their regulon specificity score (RSS). **E.** Circulating cDCs1 were FACS-sorted from peripheral blood mononuclear cells of healthy donors (n=2) and exposed to IFNβ versus PBS overnight. Left to right. FACS gating strategy, experimental scheme and lysosomal response to IFNβ assessed by Lysotracker stain by FACS.

**Supplementary Table 1. List of lysosome related genes.**

| LYSOSOMAL GENES | ENSEMBL GENE ID | PROTEIN NAME |
|---|---|---|
| Lamp2 | ENSMUSG00000016534 | Lysosome-associated membrane glycoprotein 2 |
| Lamp3 | ENSMUSG00000041247 | Lysosome-associated membrane glycoprotein 3 |
| Ctsf | ENSMUSG00000083282 | Cathepsin F |
| Ctns | ENSMUSG00000005949 | Cystinosin |
| Ctsa | ENSMUSG00000017760 | Carboxypeptidase |
| Ctsb | ENSMUSG00000021939 | Cathepsin B |
| Ctsc | ENSMUSG00000030560 | Dipeptidyl peptidase 1 |
| Ctsd | ENSMUSG00000007891 | Cathepsin D |
| Ctsf | ENSMUSG00000083282 | Cathepsin F |
| Ctsg | ENSMUSG00000040314 | Cathepsin G |
| Ctsh | ENSMUSG00000032359 | Cathepsin H |
| Ctsl | ENSMUSG00000021477 | Cathepsin L1 |
| Ctss | ENSMUSG00000038642 | Cathepsin S |
| Ctsk | ENSMUSG00000028111 | Cathepsin K |
| Lamtor1 | ENSMUSG00000030842 | Ragulator complex protein LAMTOR1 |
| Lamp1 | ENSMUSG00000031447 | Lysosome-associated membrane glycoprotein 1 |
| Gpnmb | ENSMUSG00000029816 | Transmembrane glycoprotein NMB |
| Lgmn | ENSMUSG00000021190 | Legumain |
| Vamp7 | ENSMUSG00000051412 | Vesicle-associated membrane protein 7 |
| Vamp8 | ENSMUST00000059983 | Vesicle-associated membrane protein 8 |
| Elapor1 | ENSMUST00000048012 | Endosome-lysosome associated apoptosis and autophagy regulator 1 |
| Glmp | ENSMUSG00000001418 | Glycosylated lysosomal membrane protein |
| Gga3 | ENSMUSG00000020740 | ADP-ribosylation factor-binding protein GGA3 |
| Gba1 | ENSMUSG00000028048 | Glucocerebrosidase |
| Itfg2 | ENSMUSG00000001518 | Integrin-alpha FG-GAP repeat-containing protein 2 |
| Lipa | ENSMUSG00000024781 | Lysosomal acid lipase/cholesteryl ester hydrolase |
| Lamtor2 | ENSMUSG00000028062 | Ragulator complex protein LAMTOR2 |
| Lamtor3 | ENSMUSG00000091512 | Ragulator complex protein LAMTOR3 |
| Lamtor4 | ENSMUSG00000050552 | Ragulator complex protein LAMTOR4 |
| Lamtor5 | ENSMUSG00000087260 | Ragulator complex protein LAMTOR5 |
| Laptm4a | ENSMUSG00000020585 | Lysosomal-associated transmembrane protein 4A |
| Laptm4b | ENSMUSG00000022257 | Lysosomal-associated transmembrane protein 4B |
| Laptm5 | ENSMUSG00000028581 | Lysosomal-associated transmembrane protein 5 |
| Nprl2 | ENSMUSG00000010057 | Nitrogen permease regulator 2-like protein |
| Prss16 | ENSMUSG00000006179 | Thymus-specific serine protease |
| Smpd1 | ENSMUSG00000037049 | Sphingomyelin phosphodiesterase |
| Neu1 | ENSMUSG00000007038 | Sialidase-1 |
| Lgmn | ENSMUSG00000021190 | Legumain |
| Gga2 | ENSMUSG00000030872 | ADP-ribosylation factor-binding protein GGA2 |
